# Neurotrophic Factor-Laden Acellular Chondroitin Sulfate Scaffolds Promote Chronic Functional Recovery After Severe Traumatic Brain Injury

**DOI:** 10.1101/2020.06.21.116970

**Authors:** Charles-Francois V. Latchoumane, Martha I. Betancur, Gregory A. Simchick, Min Kyoung Sun, Rameen Forghani, Christopher E. Lenear, Aws Ahmed, Ramya Mohankumar, Nivedha Balaji, Hannah D. Mason, Philip V. Holmes, Qun Zhao, Ravi V. Bellamkonda, Lohitash Karumbaiah

**Author notes:** **Correspondence:** Corresponding Author, Regenerative Bioscience Center, University of Georgia, 425 river road, Athens, GA, 30602, USA.

## Abstract

Severe traumatic brain injury (sTBI) survivors experience permanent functional disabilities due to significant volume loss and the brain’s poor capacity to regenerate. Chondroitin sulfate glycosaminoglycans (CS-GAGs) are key regulators of growth factor signaling and neural stem cell homeostasis in the brain. However, the efficacy of engineered CS (eCS) matrices in mediating structural and functional recovery after sTBI has not been investigated. We report that neurotrophic factor functionalized acellular eCS matrices implanted into the rat M1 region acutely post-sTBI, significantly enhanced cellular repair and gross motor function recovery when compared to controls, 20 weeks post-sTBI. Animals subjected to M2 region injuries followed by eCS matrix implantations, demonstrated the significant recovery of ‘reach-to-grasp’ function. This was attributed to enhanced volumetric vascularization, activity-regulated cytoskeleton (Arc) protein expression, and perilesional sensorimotor connectivity. These findings indicate that eCS matrices implanted acutely post-sTBI can support complex cellular, vascular, and neuronal circuit repair, chronically after sTBI.

## INTRODUCTION

Severe traumatic brain injuries (sTBI) caused by blunt force or penetrating trauma to the brain leads to extensive tissue loss and life-long disabilities^1^. Although sTBIs account for only 10% of the approximately 1.7 million TBI cases reported in the United States annually, they are responsible for over 90% of all TBI associated costs^1^. There are no clinical treatments to prevent cognitive impairments and tissue loss encountered after sTBI. Neural stem cells (NSCs) have long held a privileged position in neural repair strategies for their ability to mediate neuroprotective “bystander” signaling^2^. However, inadequate control over NSC differentiation and the risk of immune rejection of xenografted NSCs pose serious limitations for their clinical application^3,4^. Although autologous cell therapies could help mitigate some of these concerns^5^, issues related to poor survivability of transplanted cells^6^, and the risk of tumorigenesis in the case of human embryonic and pluripotent stem cells^7,8^, pose significant barriers to clinical success.

When compared to cell therapies, the development of acellular brain-mimetic constructs that can potentiate neurotrophic factor mediated structural and functional brain repair has received little attention. Chondroitin sulfate proteoglycans (CSPGs) are major constituents of the brain extracellular matrix (ECM) and stem cell niche^9-15^. CSPG linked sulfated CS glycosaminoglycan (CS-GAG) side chains regulate growth factor signaling^16-19^, neuronal development^20^, and neuroplasticity^21,22^. Fibroblast growth factor (FGF2) and brain-derived neurotrophic factor (BDNF) bind with high affinity to sulfated CS chains via specific sulfation motifs that act as molecular recognition sites^23^. FGF-2 is expressed by NSCs and radial glial cells found in neurogenic niches of the brain^24^, where it is known to promote neuroprotection^25^, NSC migration and proliferation^26,27^,and neurogenesis^24,26,28^. BDNF is an important mediator of neuroplasticity at all stages of brain development^29^, and is known to promote neuroprotection^30^, and functional repair of the injured brain^29,31,32^. FGF-2 and BDNF also promote angiogenesis after injury^16,33-35^, and are critical for the maintenance of oxygen perfusion and tissue viability post-injury. In recent studies, acutely implanted 3D CS-GAG scaffolds with- and without encapsulated NSCs, enhanced the efficacy of endogenous NSCs and mediated neuroprotection, four-weeks after sTBI^36^. However, the chronic functional implications of acellular, neurotrophic factor functionalized eCS matrix implants post-sTBI, has so far not been assessed.

In this study, we conducted a systematic assessment of the long-term outcomes of implanting FGF-2 and BDNF-laden sulfated eCS matrices in rats subjected to sTBI. We performed magnetic resonance imaging (MRI) phase gradient analyses of regional cerebral blood flow (rCBF) and tissue vascularization in conjunction with longitudinal behavioral performance and functional recovery, followed by detailed terminal immunohistochemical assessments of tissue-specific biomarkers. We also quantified activity-related cytoskeletal associated protein (Arc) expression and volumetric vasculature tracing in cleared brain tissue, to localize newly formed functional clusters of neurons intra- and perilesionally in sTBI rats implanted with eCS scaffolds when compared to controls.

## RESULTS

### Acutely implanted eCS matrices promote chronic neuroprotection and recovery of gross motor function

In order to assess the chronic neuroprotective effects of neurotrophic factor-laden eCS matrix implants (Sup. Fig. 1), we performed controlled cortical impact (CCI)-induced sTBI in the caudal forelimb area (CFA) in rats and implanted eCS matrices 48h post-CCI in the lesion epicenter. We performed longitudinal assessments of motor function and MRI-based quantification of tissue volume loss, and conducted terminal histological quantification of injury volume (Fig. 1A).

**Figure 1:**
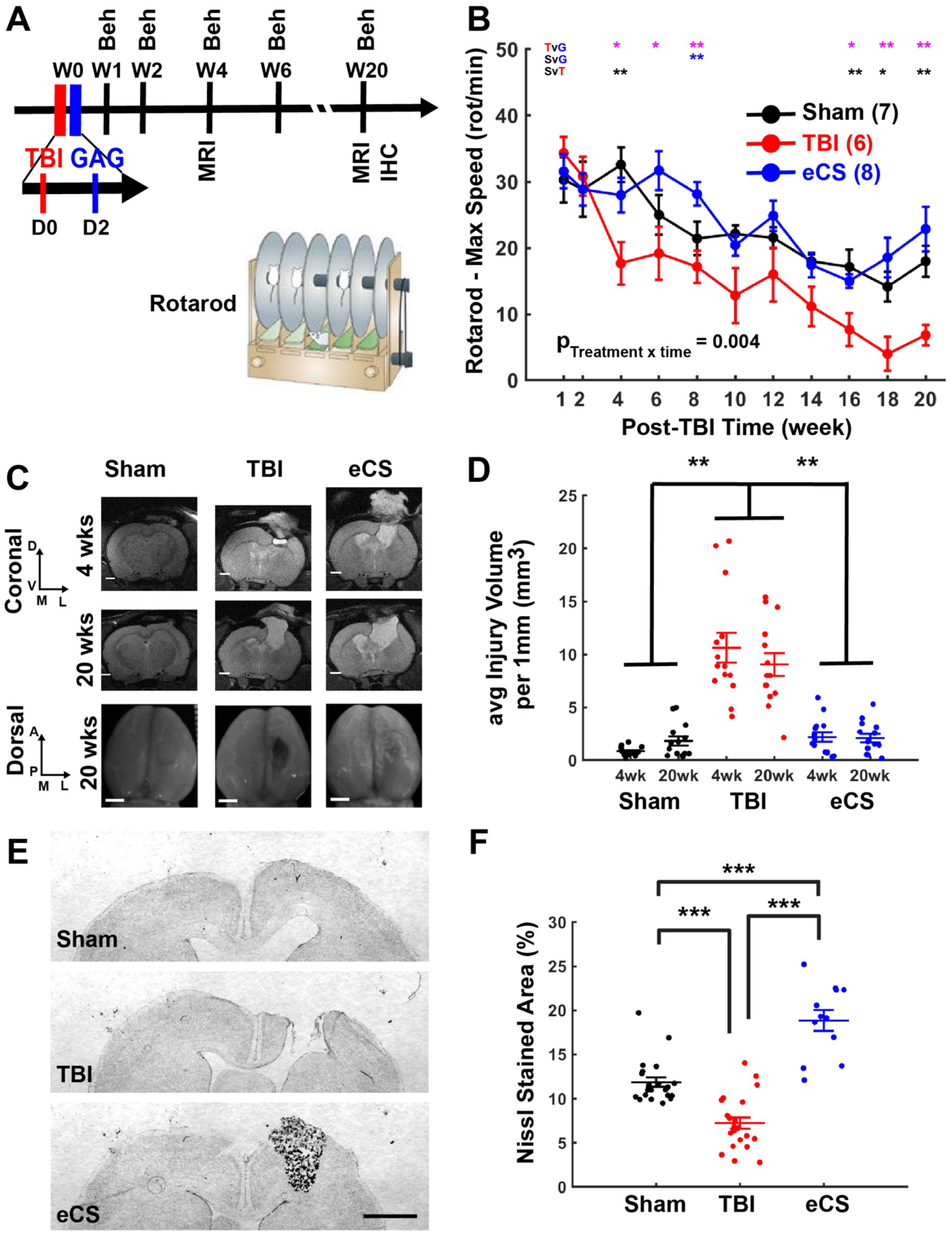
eCS implants reduce lesion volume and improve gross motor function 20 weeks post-TBI. A – Experimental schedule. All rats received a TBI at week 0 (D0). eCS implants were intracortically administered 48hour post-TBI (D2). Following one week of recovery, all rats underwent behavioral testing at week 1 and 2, and every other week therafter (4, 6, 8, 10, 12, 14, 16, 18. 20). B – The Rotarod (RR) test was used a measure of balance and motor coordination. Two-way repeated measure ANOVA; p_Treatment_ = 0.153; p_Time_ < 0.001; p_TreatmentxTime_ = 0.004. Post-hoc multiple comparison using Dunn-Sidak correction are shown above each time point for Sham vs TBI (red), Sham vs eCS(blue) and TBI vs eCS(black). C – Representative T2W MRI images (top panel) for each treatment group Sham, TBI and eCS (coronal section), scale bar: 500 μm; top view of the extracted brain (bottom panel), scale bar: 1 mm; D: dorsal, V: Ventral, M: middle, L: lateral, A: anterior and P: Posterior. D – Average injury volume quantified for each 1mm slice around the injury and based on the T2W MRI images (mm3). Two-way repeated measure ANOVA; Treatment factor: p < 0.001; Time factor: p =0.53; Treatment × Time: p = 0.0279. E – Representative Nissl stained coronal sections of rat brain for the Sham, TBI and eCS groups. Scale bar: 1 mm. F – Lesion volume quantification using Nissl stain for Sham (6 rats, 4 images per rat), TBI (6 rats, 4 images per ras) and eCS (3 rats, 4 images per rat) groups. One-way ANOVA; Treatment: p < 0.001. Post-hoc LSD *, ** and *** are for p <0.05, p < 0.01 and p <0.001, respectively. Graphs show mean ± s.e.m.

While general locomotion was found to be similar across all groups (Sup. Fig. 2A), we observed that the performance of eCS matrix implanted rats was found to be comparable to Sham controls, whereas sTBI rats demonstrated significant motor deficits (Fig. 1B and Sup. Fig. 2B).

From results of longitudinal T2 weighted MRI at 4- and 20-week time points and lesion volume analyses (Figure 1C-D), as well as terminal Nissl staining of tissue and quantification at 20 weeks post-sTBI (Fig. 1E and F), we were able to detect the significantly greater tissue atrophy in the sTBI only animals when compared to eCS matrix implanted rats and Sham controls (Fig. 1D and 1F).

These results demonstrated that eCS matrix-implanted rats exhibited tissue and motor function recovery that was comparable to Sham control group animals chronically post-sTBI.

### eCS matrices support enhanced proliferation of endogenous NSCs and neurotrophic factor expression

eCS matrices have been demonstrated to promote FGF-2-mediated neural progenitor homeostasis and neuroprotection four-weeks post-sTBI ^17,37^. Considering these previous observations, we quantified the effects of neurotrophic factor-laden eCS matrices on the chronic maintenance of proliferating NSCs in the lesion site using Ki67 and Sox-1 biomarkers (Fig. 2A & C), and CS mediated FGF-2 retention (Fig.2B,D&E). Endogenous NSC proliferation in brain tissue from eCS matrix-implanted rats was found to be comparable to Sham controls (Fig. 2C and Sup. Fig. 3A, C&D), but was significantly decreased in sTBI-only rats. Although a significant decrease in Ki67+ cells in tissue from eCS treated animals was observed when compared to Sham and TBI controls (Sup. Fig. 3A, B&E), the significantly increased percentage of Sox-1+ Ki67+ cells in eCS matrix treated animals when compared to TBI controls (Fig. 2C), suggests that NSCs do not contribute to the observed overall decrease in Ki67+ cells. These findings were further corroborated by FGF-2 expression levels (Fig. 2D), which followed a similar trend to Sox-1+ staining. CS56 antibody staining of implanted eCS matrices and endogenously produced CS-GAGs in the lesion site revealed no significant differences in CS56 staining between sTBI and eCS groups, but indicated a significantly greater CS56+ staining in both groups when compared to Sham control (Fig. 2E). Interestingly, qualitative differences in locational presence of CS56+ staining was observed throughout the lesion sites in eCS and sTBI-only animals. When compared to sTBI-only animals, which predominantly demonstrated CS56+ staining only in the lesion penumbra and indicated a complete lack of cellular presence in the lesion volume, eCS matrix-treated animals indicated a uniform presence of CS56+ staining throughout the lesion volume. The CS56+ staining of eCS matrix-implanted brain tissue indicates the potential residual presence of eCS matrix along with what appears to be endogenously produced CS-GAGs/CSPGs, 20-weeks post-TBI. These results suggest that eCS matrix implants might enhance NSC proliferation by facilitating the sustained expression and presence of FGF2 in the lesion site, 20-weeks post-sTBI.

**Figure 2:**
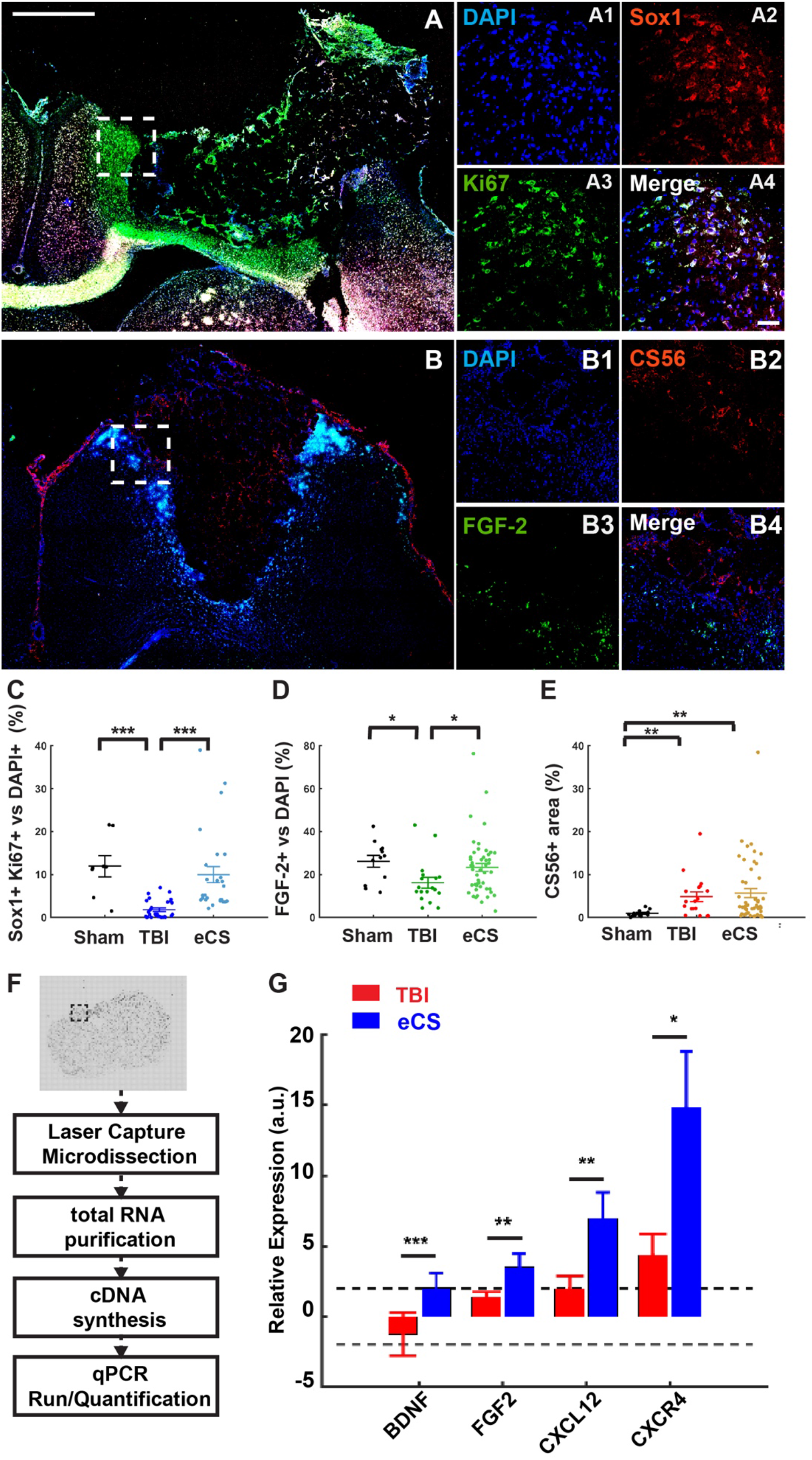
eCS implants promote NSC proliferation and neurotrophic factor expression. A – Representative tiled images of ipsilesional hemisphere (left coronal sections) of eCS rat brain tissue; scale bar = 1mm. A1-A4 – representative magnification of dashed white square shown in C for DAPI (A1), Sox1 (A2), Ki67(A3) and merged (A4); scale bar is 100 μm; B – Representative tiled images of ipsilesional hemisphere (left coronal sections) of eCS rat brain tissue; scale bar = 1mm. B1-B4 – representative magnification of dashed white square shown in A for DAPI (B1), CS56 (B2), FGF2 (B3) and merged (B4); scale bar is 100 μm; C – Co-localization Sox1+ cells with Ki67+ cells as percentage of Ki67+ cells for each treatment. Kruskal-Wallis, Treatment: p <0.001. D – Co-localization FGF-2+ cells with DAPI+ cells as percentage of DAPI+ cells for each treatment. Kruskal-Wallis, Treatment: p <0.05. E – CS56 + percentage area for each treatment. Kruskal-Wallis, Treatment: p <0.001. Post-hoc LSD Mann-Whitney U test *, **, and *** are for p<0.05, p<0.01, and p<0.001, respectively. F – At 20-week time point, brains were flash frozen and tissue was laser microdissected. Total RNA was purified and used to synthesize cDNA. qRT-PCR was run using pre-validated primers of interest for later quantification of gene expression. G – Relative expression of *BDNF, FGF2, CXCL12* and *CXCR4* transcripts in sTBI and eCS treatment groups. Two-tail t-test *, **, and *** are for p<0.05, p<0.01, and p<0.001, respectively. Graphs show mean ± s.e.m.

We used laser capture microdissection to isolate perilesional tissue labeled by rat IgG staining (Fig. 2F), along with any tissue present in the lesion cavity, in order to specifically identify regional changes in expression of *BDNF, FGF2, CXCL12*, and *CXCR4* transcripts (Fig.2G). We identified the significant and tissue-specific increase in expression levels of transcripts encoding neuroplasticity (*BDNF)* and neural progenitor proliferation (*FGF2)* factors, along with NSC homing (*CXCL12*, and *CXCR4*) transcripts in brain tissue explanted from eCS matrix-treated animals when compared to Sham and TBI controls (Fig. 2G; > 2 fold, p < 0.05). Fold differences in target gene expression in eCS matrix implanted animals was compared to that of sTBI rats after normalizing to Sham group expression levels and endogenous controls (*GADPH*, and *HPRT1*).

### eCS matrices promote chronic neurogenesis and neuroplasticity

Since cell proliferation, neuronal differentiation, and synaptic plasticity are indicators of normal brain tissue homeostasis ^38,39^, we considered the occurrence of these processes within the lesion site as measures of functional neuronal network activity, tissue maintenance, and recovery. We quantified the presence of newly formed neurons using markers for immature/migrating neuroblasts (doublecortin, DCX+) and dividing (BrdU+) neuronal cells (Fig. 3A&B). We found a significantly increased number of dividing cells in the eCS-implanted rats when compared to Sham and sTBI groups (Sup. Fig. 4A-D). A similar trend was found for DCX+ cells (Sup. Fig. 4E) and DCX+/BrdU+/DAPI+ co-labeled cells in eCS-treated animals (Fig. 3B), demonstrating a significant increase in these markers when compared to Sham and TBI groups. These results indicate that eCS matrix implants might promote the differentiation of proliferating cells into new neurons at the lesion site up to 20 weeks post-sTBI.

**Figure 3:**
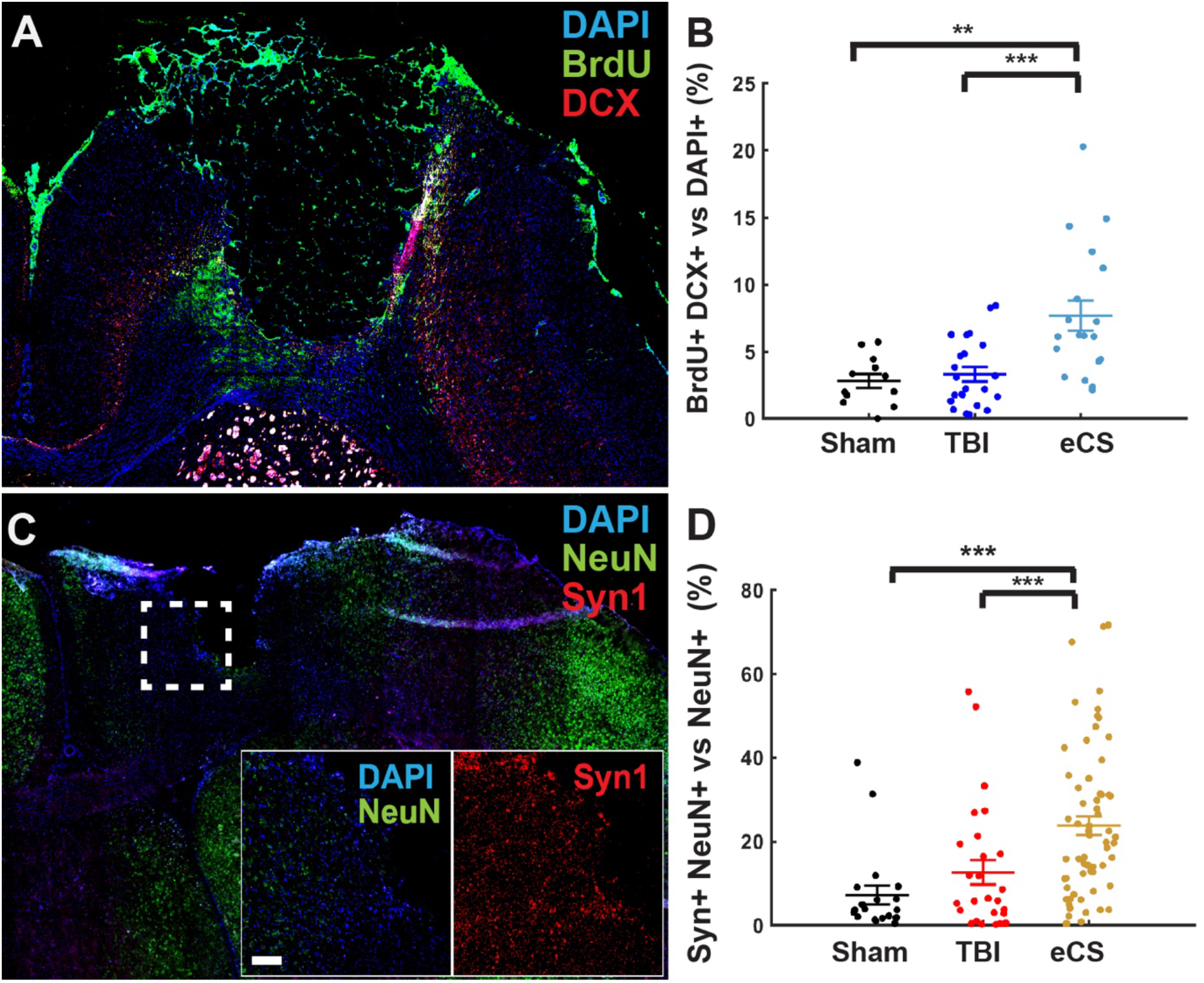
eCS implants promote neurogenesis and plasticity. A – Representative tiled images of ipsilesional hemisphere (left coronal sections) coronal sections with merged DAPI (Blue), DCX (red) and BrdU (green) staining of eCS rat brain tissue; scale bar = 1mm. B –Co-localization of DCX+ and BrdU+ cells with DAPI+ cells as percentage of DAPI+ cells for each treatments; Kruskal-Wallis, Treatment: p < 0.001. C – Representative tiled images of ipsilesional hemisphere (left coronal sections) with merged DAPI (Blue), Syn1 (red) and NeuN (green) staining of eCS rat brain tissue; scale bar = 1mm. Inset shows representative magnification of dashed white square shown as DAPI and NeuN (merged; left) and Syn1 (right); scale bar is 100 μm; D – Co-localization Syn + cells with NeuN + cells as percentage of NeuN + cells for each treatment. Kruskal-Wallis, Treatment: p < 0.001. Post-hoc LSD Mann-Whitney U test *, **, and *** are for p<0.05, p<0.01, and p<0.001, respectively. Graphs show mean ± s.e.m.

Since synaptic vesicle presence is indicative of normal functional neurons and synaptic plasticity^40^, we evaluated the potential change in synaptic plasticity at the 20-week time point using antibodies against the synaptic vesicle marker Synaptophysin-I (Syn; Fig. 3C). We observed a significant increase in Syn1+ signal in both DAPI+ (Sup. Fig. 5A-D; p = 0.014) and NeuN+ cells (Fig. 3D) in eCS matrix-implanted animals when compared to Sham and sTBI only rats.

Despite a net reduction in Olig2+ cells in eCS-implanted animals (Sup. Fig. 6C&D), we observed a significantly higher percentage of Olig2+ cells co-localized with NeuN+ cells in eCS rats when compared to those in the Sham and sTBI groups (Sup. Fig. 6A-C&E). In contrast to these findings, the co-localization of NeuN+ and Olig2+ relative to DAPI+ was found to be significantly reduced in sTBI rats (Sup. Fig. 6B&F) when compared to Sham and eCS matrix-implanted animals.

Taken together, these results suggest that despite a significant reduction in the number of neurons following TBI, eCS matrix implants promoted cell proliferation, neuronal differentiation, synaptic plasticity and potential myelination of newly formed neurons present intra- and perilesionally.

### eCS matrix implants attenuate the chronic presence of neuroinflammatory cells

Attenuated influx of neuroinflammatory cells chronically post-sTBI could prevent prolonged tissue damage and atrophy and is a marker of a favorable tissue response post-TBI ^41^. In order to mark the chronic presence of activated macrophages and reactive astrocytes that are characteristic of a neuroinflammatory cellular response to brain injury, we quantified CD68+ activated macrophages, and glial fibrillary acidic protein labeled (GFAP+) reactive astrocytes (Sup Fig. 7A-C). We observed that eCS matrix-implanted rats had similar levels of CD68+ cells (Sup Fig. 7D) as well as GFAP+ cells (Sup Fig. 7E) as Sham controls, while sTBI rats showed a significant decrease in CD68+ cells compared to Sham and eCS matrix implanted rats. Additionally, we found that CD68+ and GFAP+ cells represented a significantly lower proportion of DAPI+ cells (Sup Fig. 7F) in sTBI and eCS groups, compared to Sham controls. Although the sTBI animals demonstrated a significant reduction in activated macrophage response, this is likely due to the significant tissue loss and absence of intralesional tissue in sTBI animals when compared to Sham controls and eCS-treated animals. Overall, these results suggest the prevalence of an attenuated neuroinflammatory cellular response in sTBI and eCS animals at 20 weeks post-TBI.

### eCS matrices enhance local vascularization and global blood flow

Since inadequate vascularization is often responsible for the failure of implanted biomaterials^42^, we investigated the extent of tissue neovascularization in eCS matrix implanted animals when compared to controls, using Collagen IV (Col-IV+) and rat endothelial cell antigen (Reca+) markers (Fig.4A&B). We also used MRI phase gradient imaging of cerebral blood flow as a measure of vascular function (Fig. 4C-F).

**Figure 4:**
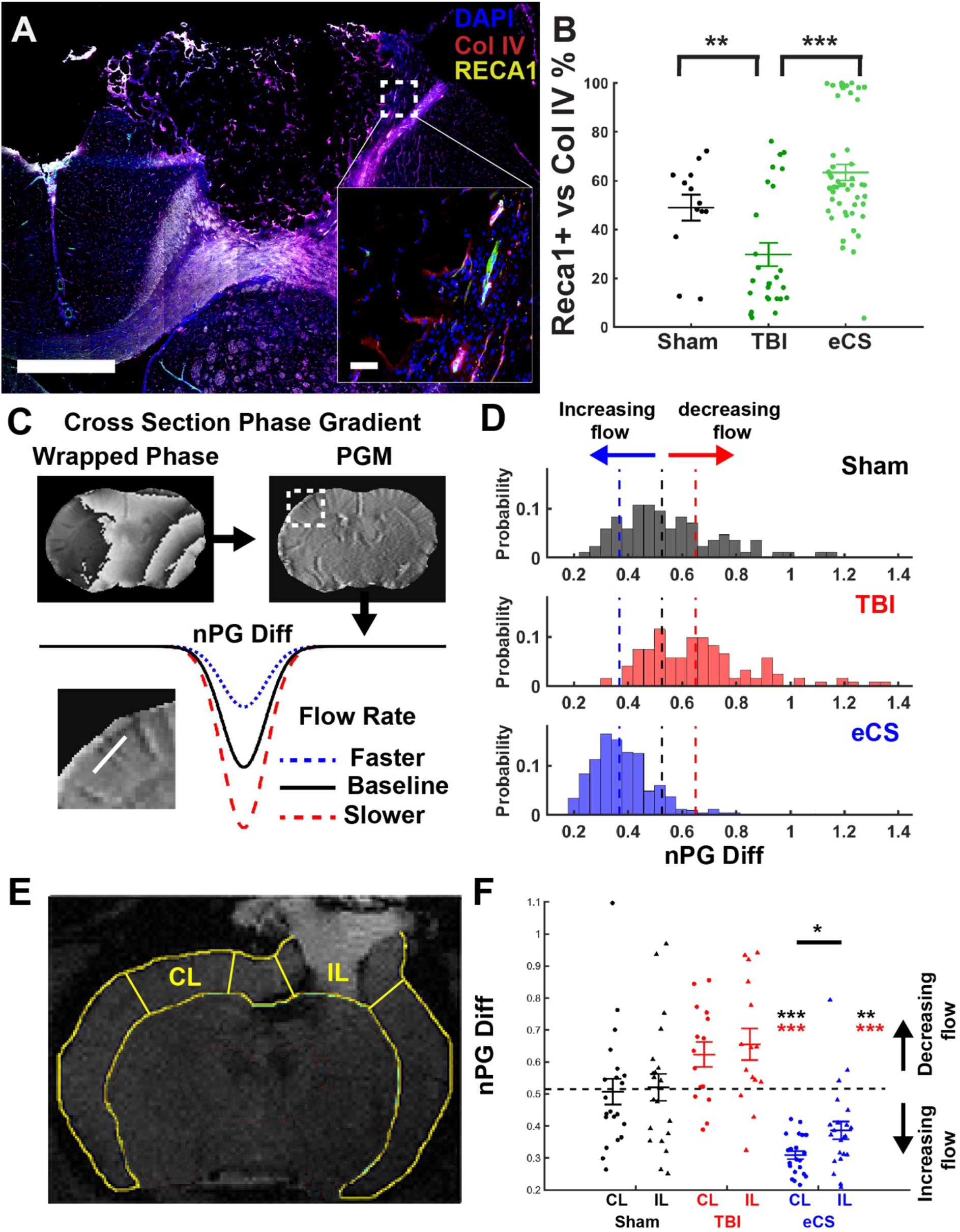
eCS promote vascularization and increased CBF 20 weeks post-TBI. A – Representative tiled image of ipsilesional hemisphere of the eCS group; scale bar = 1mm. B – Co-localization Reca1 + cells with Col IV + cells as percentage of Col IV + cells for each treatment. Kruskal-Wallis, Treatment: p <0.001. Post-hoc LSD Mann-Whitney U test *, **, and *** are for p<0.05, p<0.01, and p<0.001, respectively.4 C – Phase gradient was estimated from MRI session. First the wrapped phase map (top left panel) was used to generate phase gradient maps (PGMs, top right panel) for each slice by calculating the magnitude of the phase gradients determined in the readout and phase encoding directions (see Material and Methods for details). (bottom panel) Normalized phase gradient (nPG) differences between a vessel and the surrounding tissue were measured across all distinguishable vessels at the coronal cross section (inset, white line). Note: The rate of flow in a blood vessel is inversely proportional to the change in phase between the vessel and the surrounding tissue (see methods). D – Distribution of nPG for all identified blood vessel in each treatment groups Sham (n = 84), TBI (n = 121) and eCS (n = 432). Comparison using the glass delta measure effect size and 10,000 bootstrapping. Dashed line represent the median nPG for Sham (black), TBI (red) and eCS (blue). One-way ANOVA, F(2) = 230.56, p = 8.57e-51 / measure effect size p < 0.001. E – Representative segmentation of MRI coronal section in sub-region for regional and hemispheric comparison in blood flow. IL: ipsilesional, CL: contralesional. F – nPG for the perilesional region of interest and matching region of interest in the contralateral hemisphere. Two-way ANOVA, Treatment: p <0.0001. Post-hoc one-sample t-test; Post-hoc LSD *, **, and *** are for p<0.05, p<0.01, and p<0.001, respectively. Graph shows mean ± s.e.m.

We found a significant increase in Reca-1+ (Sup. Fig. 8D) and Col-IV+ (Sup. Fig. 8E) percentage area expression in the eCS matrix-treated animals compared to both Sham and sTBI controls. Notably, Reca-1 and Col-IV co-localization was detected to be ~60% in Sham and eCS matrix-treated groups, which was significantly enhanced when compared to the sTBI group (Fig. 4B).

Since a significant reduction of cerebral blood flow (CBF) chronically is linked to cognitive dysfunction and poor prognosis in humans with sTBI^43-47^, and is also associated with the lack of neuronal activity and loss of neuronal volume in humans and rats^48,49^, we assessed CBF using MRI-based normalized phase gradient mapping of identified blood vessels in the TBI lesion (nPG; Fig. 4C). We observed a significantly enhanced overall cerebral blood flow (CBF) in eCS matrix-implanted rats (Fig. 4D and Sup. Fig. 9; estimation statistics) when compared to Sham animals, while sTBI control animals showed a nonsignificant decrease in measured CBF. Using ipsi- and contralesional region of interest measurements (Fig. 4E), we detected a specific increase in cortical CBF in eCS matrix implanted rats with a significantly higher CBF detected in the contra- vs ipsilesional side (Fig. 4F). Taken together, these results indicate that eCS matrix implanted rats demonstrate chronically enhanced neovascularization perilesionally, along with significantly enhanced CBF both contra- and ipsilesionally when compared to TBI only controls.

### eCS matrices promote chronic forelimb-specific functional recovery and activation of activity-regulated cytoskeleton-associated protein (Arc) in RFA

We used a skilled reach task in order to investigate the circuit-specific implications of the neuroprotective, neurogenic, and angiogenic attributes of eCS matrix implants following sTBI. The skilled reach task was used as a forelimb-specific assessment of motor recovery followed immediately by terminal volumetric imaging of activity-regulated cytoskeleton-associated protein (Arc+). This made it possible to assess task-specific responses following a lesion of the M2 – reach-to-grasp region in the rostral forelimb area (RFA).

We found that eCS matrix-implanted rats that trained for 2 weeks on the skilled reach task (Fig. 5A), demonstrated reach-to-grasp performance recovery from week 2 that was comparable to Sham animals, and that persisted up to week 8 (Fig. 5B and Sup Fig. 10). sTBI control rats showed significant forelimb functional deficits throughout the 8 weeks of testing, with a transient improvement that lasted about 3 weeks (week 4 through 6) before significantly worsening at week 8. The quantification of ipsi- and contralesional Arc+ neurons (Arc+/NeuN+ co-expression; Fig. 5C) immediately post reach and grasp activity, showed strong activation within anterior and posterior RFA regions and low contralateral signal in Sham animals (Fig. 5D). sTBI control rats, in comparison, showed low to no ipsilesional RFA activity with notable contralesional activation when compared to eCS matrix-implanted rats, which demonstrated Arc+ signal within the RFA lesion in addition to contralesional activation.

**Figure 5:**
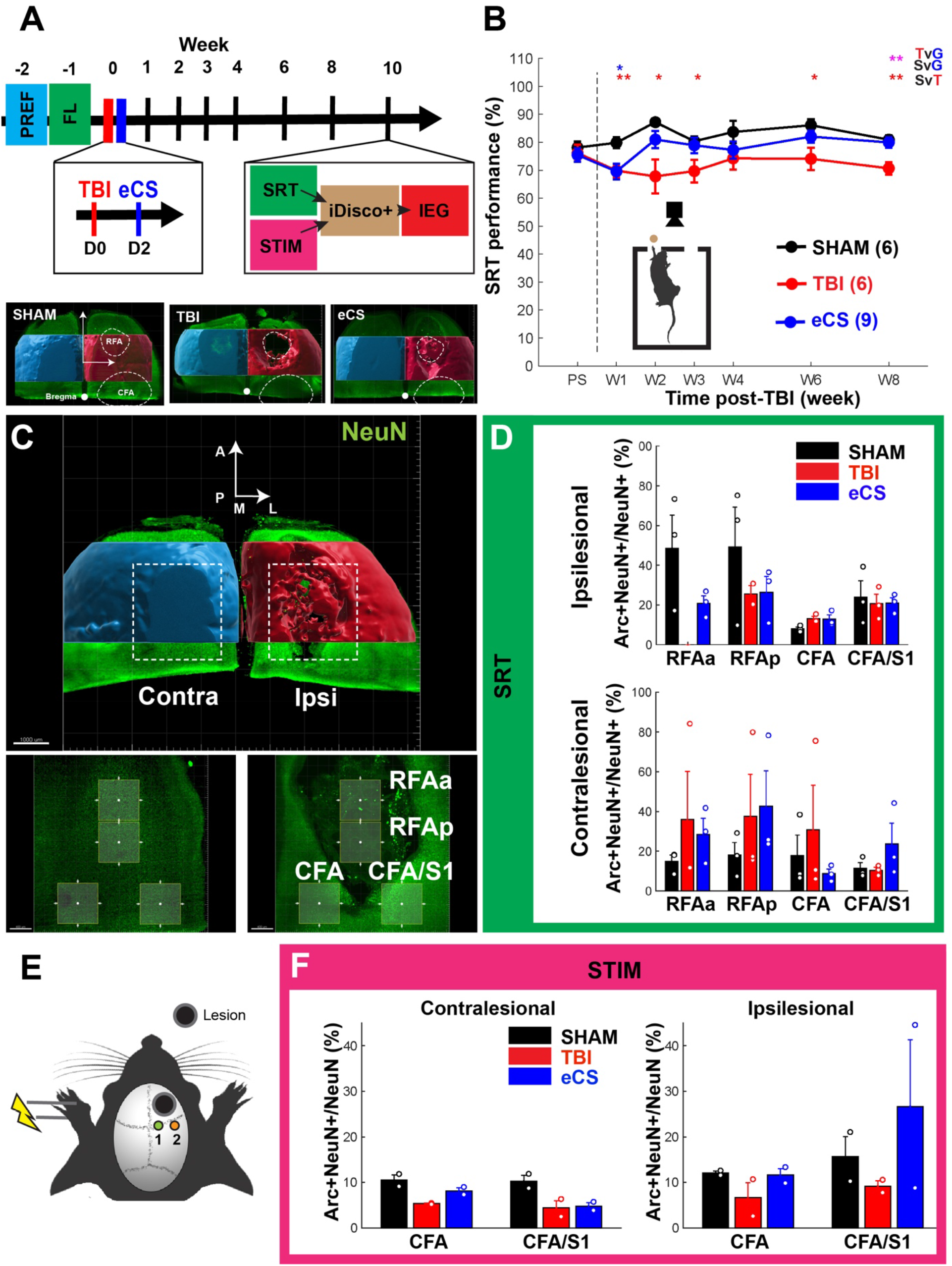
eCS implanted animals demonstrate enhanced recovery of Reach-to-Grasp-specific motor function 8 weeks post-TBI. A – Experimental schedule of skilled reach task. (Top panel) All rats received training for the skilled reach task starting from a preference training followed by a forced left limb training. A CCI at the rostral forelimb area (RFA) was performed followed by eCS implants (48h post-sTBI). Rats were tested weekly/bi-weekly on the skilled reach task. At week 10, Rats were terminally sacrifices and brain processed for immediate early gene expression following either an intensive skilled reach task assay or contralesional paw stimulation/brain recordings. (Bottom panel) Representative volumetric segmentation of lesion from iDisco+ cleared and NeuN+ stained brains. Representative images showing tissue presence for Sham (left), TBI (middle) and eCS (right) groups using surface mapping in Imaris. The contralesional and ipsilesional hemispheres are color coded in blue and red, respectively. B – Overall performance for the skilled reach task in percentage of the Sham (n = 6), TBI (n = 6) and eCS (n = 9) groups post-TBI and treatment. The SRT performance score is calculated from the average between success hit (%), left-limb usage (%) and efficiency (%). Repeated measure ANOVA, Time: p <0.05, Group: p <0.01, Time × Group: p = 0.1306; Post-hoc LSD * and ** indicate a p<0.05 and p<0.01, respectively. The color of stars indicates statistical test between the following pairs: Sham vs TBI (SvT, red), Sham vs eCS (SvG, Blue) and TBI vs eCS (TvG, magenta). RFAa: RFA anterior; RFAp: RFA posterior. SRT: Skilled reach task. C – Region of Interest (ROI) definition for the analysis of Arc+ NeuN+ colocalization for reach-to-grasp function mapping. The ipsilesional and contrlaesional sides are marked in red and blue, respectively. All rats were sacrificed at week 10 post-sTBI, 30 min post SRT assay, transcardially perfused, and brain tissue processed for iDisco+ immunostaining. Imaris quantification of lightsheet images was used to determine the percentage of Arc+ expressing NeuN+ cells within each ROI. The contralesional and ipsilesional hemispheres are color coded in blue and red, respectively. D – Ipsilesional (top panel) and contralesional (bottom panel) quantification of Arc+ NeuN+ colocalized cells post-SRT assay reveals RFA specific activation of forelimb circuitry in Sham as well as in eCS rats. TBI rats showed absence of cell or activity in the RFAa. Both TBI and eCS rats showed increased perilesional and contralesional activity 10 weeks post-TBI. Sham (n = 3), TBI (n = 3) and eCS (n=3). Line plots and bar graphs show mean ± s.e.m. E – Experimental setup for the paw stimulation combined with laminar electrophysiological recording. Rats were sacrificed after stimulation-recording to quantify neuronal activity-induced Arc expression in somatosensory regions. F – Contralesional (left panel) and ipsilesional (right panel) quantification of Arc+ NeuN+ colocalized cells post-STIM assay for Sham (n = 2), TBI (n = 2) and eCS (n = 2). Bar graph show mean+/-s.e.m.. Line plots and bar graphs show mean ± s.e.m.

### Perilesional laminar recordings of multi-unit activity reveal the preservation of sensorimotor responses 10 weeks post-TBI in eCS matrix implanted rats

We stimulated the contralesional paw using low intensity electrical pulses to further investigate whether eCS matrix implants directly facilitated recovery of sensorimotor responses (Fig. 5E; Sup. Fig. 11A). We evaluated the spontaneous and evoked (Sup. Fig. 11B and C) electrophysiological responses from the intact M1 and M1/S1 regions using a 32-channel laminar electrode (Sup. Fig. 11D). We observed that following sTBI, a rapid response to electrical stimulation of the left paw reduced in occurrence rate and increased in time jittering in all cortical layers and in the two recording positions CFA and CFA/S1 (Sup. Fig. 11E). Notably, the late response recorded in the CFA region in response to paw stimulation showed an increased delay in sTBI animals (Sup. Fig. 11E left; mean jitter: 36.5 msec), which was found to be faster in eCS matrix implanted rats (mean jitter: 15 msec). This effect was not detected in CFA/S1 region (Sup. Fig. 11E right). We then looked at the region activation pattern in stimulated/recorded rats using Arc+ labeling as in Fig. 5D (Fig. 5F). We found that the level of activity recorded in CFA and CFA/S1 returned to Sham levels in eCS rats, while sTBI control rats showed a lower activity level following contralateral paw stimulation. The sTBI control rats also showed a sustained post-stimulation activity registered during non-stimulation recording periods when compared to the pre-stimulation epochs (Sup. Fig.12), which was not observed in Sham and eCS matrix-implanted rats. Consistently with forelimb-specific circuit activation, we found that paw stimulation induced activation of the CFA/S1 area and associated motor area CFA with TBI and eCS matrix implanted rats showing a marked reduction and increased activation, respectively. These results indicate that the response and activation of the perilesional circuitry associated with sensorimotor function of forelimb was reduced in TBI rats and partially recovered in eCS matrix-implanted rats.

### RFA-specific re-vascularization explains neuronal presence and behavioral performance

We performed brain tissue clearing to specifically investigate whether vascular architecture could reveal a stronger correspondence between vasculature features (Fig. 6), neuronal presence, and behavioral performance.

**Figure 6:**
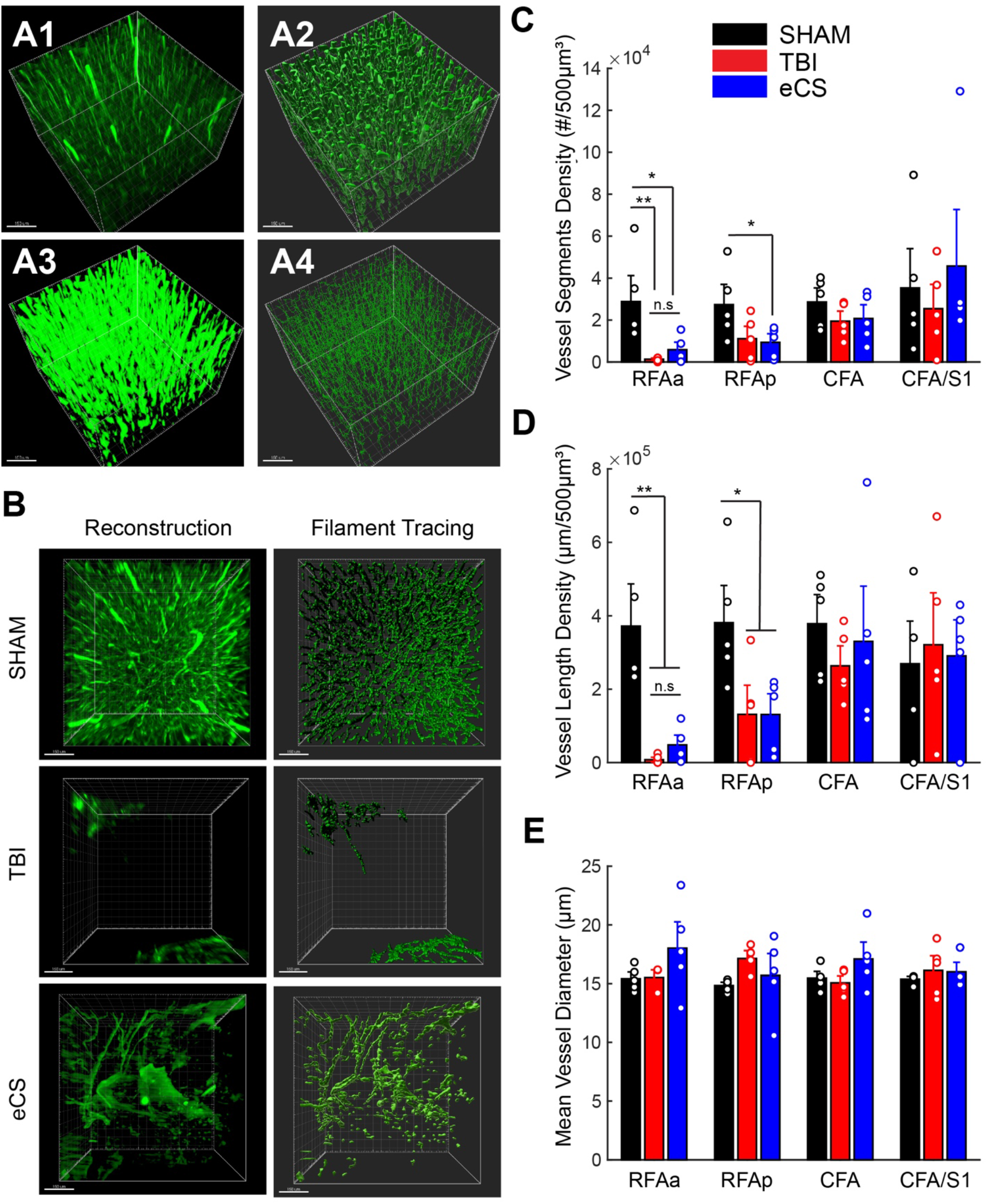
eCS implants promote changes in intra and perilesional vascular architecture 10 weeks post-TBI. A – Image processing step for vasculature tracing. Using Imaris software on volumetric imaging of iDisco+ cleared brains, we first used reconstructed volumetric images from Reca1+ staining (green – A1), performed surface mapping (A2), surface-based masking and Gaussian filtering (A3) and finally filament tracing (A4). Representative vascularization in eCS implanted region; original Reca1+ images (left) and vasculature traced ROIs (right) B – Representative images of Sham (Top panel), TBI (middle panel) and eCS (bottom panel) rat brain tissue for the reconstructed original volume image (left) and vasculature tracing (right). C – Quantification of blood vessel segment density per 500 μm^3^ for Sham (n = 15), TBI (n = 15), eCS (n = 15) groups in all four ROIs as in figure 6. D – Quantification of blood vessel length density per μm/500 μm^3^ for Sham (n = 15), TBI (n = 15), eCS (n = 15) groups in all four ROIs as in figure 6. E – Quantification of mean blood vessel diameter (Volume: 500 μm^3^) for Sham (n = 15), TBI (n = 15), eCS (n = 15) groups in all four ROIs as in figure 6. Post-hoc LSD Mann-Whitney U test *, ** and *** indicate p < 0.05, p < 0.01 and p < 0.001, respectively. Bar graphs show mean ± s.e.m. RFAa: RFA anterior; RFAp: RFA posterior/perilesional; Graphs show mean ± s.e.m.

Following Imaris-based vasculature tracing (Fig. 6A and B), we observed that the anterior RFA region in eCS rats showed an increased vessel segment density (Fig. 6C) and cumulative vessel length (Fig. 6D) when compared to TBI control rats, whereas both TBI and eCS matrix-implanted animals demonstrated a marked reduction in these features compared to Sham. Notably, only 3 out of 5 rats showed tissue presence in the RFAa ROI in TBI controls, whereas all animals in Sham and eCS matrix-implanted groups showed cellular and tissue presence. Tissue clearing and volumetric imaging revealed a larger propensity of tissue preservation in eCS matrix-implanted rats compared to sTBI controls (Sup. Fig. 13), which is consistent with lesion volume analysis obtained in Fig. 1. We also observed that the posterior RFA showed a decreased density of vasculature in both sTBI and eCS matrix-implanted animals.

Interestingly, we found that the mean vessel diameters of vasculature in sTBI controls and eCS matrix-implanted animals were not significantly different from those in Sham rats (Fig. 6E). However, consistently with the increased CBF observed in eCS matrix implanted animals (Fig. 4 – 20 weeks), the distribution of both vessel length density and diameter in RFAa were enhanced in eCS matrix-implanted rats when compared to Sham and TBI controls, as shown in Q-Q plots (Sup. Fig. 14A and B).

Importantly, we found a significant correlation between NeuN+ cells and vessel density (Sup. Fig. 15A), and NeuN+ cells and forelimb performance (Sup. Fig. 15B). The correlation between vessel density and skilled-reach task performance also returns a strong Pearson’s R-value (Sup. Fig. 15C), although not significant. Taken together, these results indicate that eCS matrix implants promoted vascularization intralesionally (RFAa), which strongly correlated with neuronal presence and forelimb performance in rats.

## DISCUSSION

Parenchymal volume loss of brain tissue is highly correlated with the level of TBI severity in humans, with patients sustaining severe lesions experiencing significantly greater volume loss when compared to those who sustained mild TBIs^50^. As a result, functional losses are inevitable and neuroplastic changes are limited, leaving patients with chronic and debilitating impairments^1^. Studies on the intracortical implantation of brain-mimetic CS and hyaluronic acid GAG scaffolds after TBI and stroke have demonstrated the ability to mediate complex structural and functional repair of brain tissue^37,51,52^. The acute implantation of acellular eCS scaffolds that are functionally diverse and compositionally similar to brain ECM, could present a rational approach to mitigating the significant volume and functional losses encountered chronically after sTBI.

Using a materials design strategy that exploits the native functionality of CS, we demonstrate that neurotrophic factor-laden eCS matrix implants were neuroprotective, neurogenic, and significantly enhanced the peri- and intralesional presence of newly formed blood vessels chronically after sTBI when compared to controls. The role of a global reduction of CBF and hypoperfusion in chronic dysfunction and poor prognosis post-sTBI is well-established^44-48^. Our observations correlating enhanced vascular density to neuronal presence and reach-to-grasp function recovery, suggests that eCS matrix implants orchestrate complex vascular repair that directly contributes to neuronal function and task-specific recovery chronically post-sTBI. These effects are likely mediated by the enhanced presence and signaling of FGF-2 and BDNF, which is potentiated by CS binding of these factors as demonstrated previously^17,51^.We nonetheless interpret the implications of observed rCBF increase cautiously in the absence of comparative real-time cerebral oxygenation studies.

The compensatory reorganization of the motor cortex has been observed in regions spanning the perilesional circuitry and the contralateral hemisphere post-TBI^53-56^. In our studies, the volumetric quantification of Arc+ activated neuronal populations following left-limb usage confirmed enhanced activity in both the perilesional and contralateral regions in sTBI-only rats compared to Sham controls. eCS matrix-implanted rats showed a pattern of functional activity similar to sTBI-only rats, with the added activity of intralesional neurons within the implant. Our results also showed a close correspondence between Arc-dependent activation patterns, and both electrophysiological response and recovery of forelimb-specific sensorimotor functions. This evidence suggests that intracortical eCS implants mediated the reorganization of neuronal circuitry leading to chronic recovery of reach-to-grasp function post-sTBI.

In summary, our results demonstrate that rationally designed, brain-mimetic, acellular eCS implants possess native structure-function attributes required to mediate chronic tissue level repair and functional recovery post-sTBI. This study opens up new avenues for the design and application of synthetic sulfated GAG constructs for brain tissue repair, and informs future investigations of neuronal connectivity and electrophysiological responsiveness between the eCS matrix implanted tissue and perilesional cortical columns.

## Author Contributions

C.-F.L, M.B, and LK designed experiments.

C.-F. L., M.B, H.M., R.F. and M.S. performed all animal behavior and surgeries.

N.B. and L.K. performed LCM and qRT-PCR experiments.

M.B performed MRI imaging and G.S. performed phase gradient analysis.

C.-F.L., M.S. and A.A performed the tissue clearing, volumetric imaging and volumetric quantification.

C.-F.L., M.S., R. F., N.B and C.E.L. performed all samples cryosectioning, staining and imaging.

C.-F.L., R.M. and N.B performed IHC-based quantification, behavior analysis, qRT-PCR analysis and statistical analysis.

C.-F. L. and L.K. wrote the manuscript and C.-F.L., Q. Z., P. H., R.V.B and L.K. edited the manuscript.

## DATA AVAILABILITY STATEMENT

Data presented and analyzed in this manuscript are available upon request from

## MATERIALS AND METHODS

### A – Animal procedures and preparation of eCS matrix

#### 1. Animals

Sprague-Dawley rats were obtained for the 20 weeks (n = 33, Harlan Laboratories, 7 weeks old; males only) and the 10 weeks experiments (n = 21, Charles River Laboratories, 3 weeks old; males only), and randomly assigned to each treatment: control craniotomy with no injury (SHAM group; 20 weeks: n = 9; 10 weeks: n = 6), Controlled Cortical Impact (CCI; TBI group; 20 weeks: n = 9; 10 weeks: n = 6), and CCI implanted with eCS scaffolds (eCS group; 20 weeks: n = 15; 10 weeks: n = 9). A custom-designed controlled cortical impactor delivered the desired impact to the M1 motor cortex (20 weeks: Caudal Forelimb Area, CFA; AP: 0.5 mm; ML: 0.5 mm, relatively to bregma; 10 weeks: Rostral Forelimb Area, RFA; AP: 3.0 mm; ML: 2.5 mm, relatively to bregma) of the eCS matrix implanted animals and TBI-only controls.

All rats were single housed with *ad libitum* access to food and water and following a reverse 12h-light cycle (light OFF 7:00-19:00, Light ON 19:00-7:00) in a room maintained at 70% humidity and 23°C. All procedures on animals were approved by the Institutional Animal Care and Use Committee (IACUC), and protocols were performed in accordance with the Guide for the Care and Use of Laboratory Animals published by the National Institution of Health (NIH).

#### 2. Surgical procedures

Prior to CCI injury, each animal was anesthetized with 5% isoflurane and buprenorphine was injected subcutaneously (0.3mg/ml, 0.05 mL/300g; Henry Schein). Animals were placed on a stereotaxic frame attached to a temperature controlled heating pad (37°C) with their scalp shaved and sanitized (70% EtOH and 3% Povidone-Iodine). Following, a sagittal incision, the periosteum was cleaned using Etch-gel (Phosphoric acid Etching; Henry Schein), and a craniotomy was performed using a 5-mm-diameter trephine bur fitted to an electronic drill. A 3-mm CCI tip was fitted onto the pneumatic piston and positioned in contact with the surface of the dura (fully extended position), and then retracted to adjust for an impact depth of 2mm. A severe CCI injury was induced by programming the piston speed to 2.25 m/s and a dwell time of 250 ms, resulting in an initial 3-mm-diameter injury with a depth of 2 mm. Absorbable gelatin (Gel foam ^®^, Pfizer pharmaceutical) was applied to the injury site, and sterile cotton swabs were used to remove excess blood. The Gel foam was then removed, and the injury site was covered completely with a layer of 1% sterile SeaKem (Lonza) agarose. Skin flaps were sutured together, closing the wound. Triple antibiotic cream was applied on the sutured skin.

#### 3. Preparation and intralesional delivery of eCS matrix

Injured animals were randomly selected to receive intralesional injections of eCS matrix functionalized with the neurotrophic factors fibroblast growth factor (FGF-2; 50μg/mL; R&D Systems; 233FB025CF) and brain-derived neurotrophic factor (BDNF; 50μg/mL; R&D Systems; 248BDB010CF), 48h post-sTBI. The eCS matrices were prepared using previously described methods ^17,36^ (22 uL: 8 μL of 10% methacrylated-chondroitin-sulfate-A-glycosaminoglycan; 7.58 μL neurobasal media; 4 μL of 1% photocrosslinker Irgacure-2959, Sigma-Aldrich; 1.21 μL of FGF-2; 1.21 μL of BDNF), loaded in a 50 μL Hamilton Syringe (fitted blunted 21 Gage tight lock stainless steel needle) and photo-crosslinked for 1min just prior to injection. Injection was performed following reopening of the scalp following the above described procedure, removing the 1% agarose and positioning the syringe (angle of 32°, depth: 2mm; speed: 2uL per min for 10 min). Buprenorphine was administered (0.3mg/ml, 0.05 mL/300g; Henry Schein) and the skin was sutured back.

### B - Behavioral assessment of functional recovery

Chronic functional recovery was assessed using longitudinal gross motor recovery assays including the beam-walk test (BW)^57^, rotarod test (RTT)^58^ and open field test (OFT) for time points corresponding to week 1, 2, 4, 6, 8, 10, 12, 14, 16, 18, and 20 post-TBI. Chronic recovery of forelimb-specific function was assessed using the skilled reach task (SRT) following preference and forced-side training at 1, 2, 4, 6, and 8 weeks post-TBI. All tasks were performed during the dark cycle in a red-light-lit room held at constant 70% humidity and 23°C.

### D - Magnetic Resonance Imaging (MRI)

#### 1. MR imaging procedure

A 7T Varian Magnex MRI scanner and a 35mm-dualtuned surface coil were used to perform MRI of the brain in order to investigate the chronic (20-weeks) effects of TBI. All animals were placed directly under isoflurane (1.5%) anesthesia on the holder of the animal-tube assembly where a catheter was added to the tail of each rat intravenously with a line prefilled with 0.2ml heparin flush attached. The animal’s head was securely positioned directly under the 20mm-dualtuned surface coil, and the animal-tube assembly was placed inside the 210 mm horizontal bore of the small animal scanner. For each imaging sequence, eight coronal brain slices with 1 mm thickness and zero gap between slices were acquired using a field of 40×40 mm and a matrix size of 256×256. The following two imaging sequences were acquired for each animal: (1) T_2_-weighted fast spin echo (T2WI): TR = 2.0 s, TE = 32 ms, and 4 averages requiring an acquisition time of 4 min and 20 sec and (2) T_2_*-weighted multi-gradient-echo (MGEMS): TR=600ms, 8 echoes (TE=10-45ms in 5ms increments), FA=25°, and 4 averages requiring an acquisition time of 10 min and 14 sec. Therefore, the total acquisition time for each animal was ~15 min.

#### 2. Injury volume determination

As T2WI images revealed the largest areas of injury and inflammation, T2WI images were used to design a region of interest (ROI) library for each animal’s injured cortex using ImageJ (National Institute of Health, MD) to manually segment the injured area.

#### 3. Phase-gradient mapping

The rate of flow in a blood vessel is inversely proportional to the change in phase between the vessel and the surrounding tissue ^59^. However, during MR acquisition, phase measurements are only recorded in the range of ±π, and any values that extend outside of this range get “wrapped”, or forced into this range, making calculating the change in phase difficult in areas where wrapping occurs. Since the change in phase is proportional to the change in the phase gradient, phase gradient maps (PGMs) can be used to avoid this wrapping problem ^60^.

Changes in rCBF between groups was determined using the T2*-weighted MGEMS data using the following procedure ^61^. Phase gradient maps (PGMs) were generated for each slice by calculating the magnitude of the phase gradients determined in the readout and phase encoding directions ^60^. From the PGMs, phase gradient differences between a vessel and the surrounding tissue were measured across all distinguishable vessels located within the cortex, sub-cortical and basal forebrain areas, and thalamus. The third echo (TE=20ms) was used for all measurements as this echo produced the best contrast for distinguishing vessels within the PGMs. To account for systematic phase variations in the MRI acquisition across rats (differences in coil tuning, gradient shimming, etc.), measurements were normalized using the phase gradient difference measured across the third ventricle for each rat. Averages and standard deviations of all normalized phase gradient differences were calculated for all vessels located on both the contralateral and ipsilateral side of the brain to the injury for each region and group.

### E – Brain tissue preparation and immunohistochemistry

#### 1. Tissue collection at 20-week time point

20 weeks post-injury, animals were heavily sedated using ketamine (65mg/kg) and transcardially perfused with 250 mL 0.1 phosphate-buffered saline (PBS, pH 7.4) followed by 250 mL 4% paraformaldehyde solution. Brains were then extracted and cut in half (transversal cut), 7mm from the apex of the frontal cortex, flash frozen in liquid nitrogen and stored at −80°C. Coronal sections were obtained using a Leica Cryostat, starting from the center of the injury and moving caudally. A total of 48, 15-μm-thick, sections were collected per animal and distributed evenly over 10 charged slides and 2 membrane slides (4 sections per slide). 2 slides per animal were placed in 4% paraformaldehyde containing 4M sucrose, and stained with 0.1% cresyl violet solution to label Nissl bodies in neurons. The remaining 8 charged slides were used for immunofluorescence assays, and the 2 membrane slides were used for qRT-PCR. For immunostaining, slides were dried and placed in 4% paraformaldehyde solution containing 4M sucrose, assigned to one of the primary antibody groups and incubated overnight at 4°C (Table 1). Slides were then incubated with blocking buffer consisting of 1:220 dilutions of appropriate secondary antibodies (Table 2), counterstained with cell nuclear stain, NucBlue ^®^ (Life Technologies, NY), mounted with Fluormount-G (Southern Biotech, AL), sealed with coverslips and stored at −20°C until imaged.

**Table 1:** Primary Antibody panel for immunostaining

| Purpose | Antibody | Manufacturer | Cat# | Host | Dilution | Usage |
| --- | --- | --- | --- | --- | --- | --- |
| Neuronal<br>Presence/marker | NeuN | EMD<br>Millipore | MAB377 | Mouse | 1:500 | Slices |
| Neuronal<br>Presence/marker | NeuN | EMD<br>Millipore | ABN91 | Chicken | 1:500 | iDisco+ |
| Neuronal<br>Proliferation | Doublecortin<br>(DCX) | Abcam | ab18723 | Rabbit | 1:500 | Slices |
| Myelination | Olig2 | Abcam | ab109186 | Rabbit | 1:100 | Slices |
| Neural<br>Progenitor | Sox-1 | Abcam | ab87775 | Rabbit | 1:1000 | Slices |
|  | Ki67 | Santa Cruz | sc-7846 | Goat | 1:500 | Slices |
| Neuroinflammation | GFAP | DAKO | z0334 | Rabbit | 1:1000 | Slices |
|  | CD68 | Bio-Rad | mca341R | Mouse | 1:500 | Slices |
| Growth Factor<br>presence | FGF | Abcam | ab8880 | Rabbit | 1:200 | Slices |
| Chondroitin<br>Sulfate GAG<br>presence | CS56 | Sigma-Aldrich | C8035 | Mouse | 1:200 | Slices |
| Plasticity | Synapsin-1 | Abcam | ab14692 | Rabbit | 1:500 | Slices |
| Neuronal<br>Activation<br>Marker | Arc | Synaptic<br>Systems | 156 003 | Rabbit | 1:1000 | Slices<br>/iDisco+ |
| Vasculature | RECA | BioRad | MCA970R | Mouse | 1:500 | Slices<br>/iDisco+ |
|  | Col IV | Abcam | Ab19808 | Rabbit | 1:500 | Slices |

**Table 2:** Secondary Antibody panel

| Alexa-Fluor | Host | Reactivity | Isotype | Catalog # | Company |
| --- | --- | --- | --- | --- | --- |
| 488 | Goat | Mouse | IgG (H+L) | A11006 | Life Technology |
| 488 | Goat | Chicken | IgY (H+L) | A11039 | Life Technology |
| 647 | Goat | Rabbit | IgG (H+L) | A21244 | Life Technology |
| 488 | Donkey | Goat | IgG (H+L) | A11055 | Life Technology |
| 594 | Goat | Mouse | IgG (H+L) | A11005 | Life Technology |

#### 2. Laser capture microdissection (LCM)

LCM was performed on two membrane slides per animal as previously described^62^. In brief, brain tissue sections were placed on a cold block and fixed for 2 minutes in ice-cold 75% ethanol solution then rinsed thrice with ice-cold nuclease free PBS (pH 7.4, Life Technologies). Brain sections were stained in a 1:220 dilution of goat anti-rat IgG + 500 U RNase inhibitor (Life Technologies). After 1-hour exposure, sections were rinsed three times with nuclease free PBS and dehydrated sequentially with 75%, 95%, 100% ethanol for 15 seconds each. Sections were placed on the ice block to dry for 5 minutes prior to LCM using a laser microdissection microscope (Leica).

#### 3. qRT-PCR

Total RNA was extracted from LCM processed tissue using a picopure RNA isolation kit (ThermoFisher). and quantified using a nanodrop analyzer (ThermoFisher). 100 ng equivalent of pooled total RNA (5 slices per animal; 7 rats) was used to synthesize cDNA using an RT First Strand Kit (Qiagen). Pre-validated primers targeting rat *CXCR4, CXCL12, BDNF, FGF-2*, along with the housekeeping genes *HPRT1* and *GAPDH* (Qiagen) were used in qRT-PCR assays for 40 cycles in three steps: 95°C for 10 minutes, 40 cycles of 95 °C for 15 seconds, and 60 °C for 1 minute. Melting curve analysis was conducted to validate amplification specificity. All assays were conducted in triplicate on a 7900 HT instrument (Applied Biosystems) and gene expression levels normalized to endogenous gene controls plotted as relative units using the ΔΔCt method. Gene expression was analyzed with an absolute fold difference greater than 2. Gene expression was normalized against the average expression of SHAM controls.

#### 4. Tissue collection and preparation for task and stimulation specific activation of immediate early genes – 10 weeks

For the visualization of functional activity in the whole brain, we used antibody staining against the immediate early gene (IEG) encoded actin-regulated cytoskeleton-associated protein (Arc). Two groups of rats were used at 10 weeks post-TBI. The first cohort of rats performed a continuous 40 min of SRT (specific left forelimb activation with high repetition; n = 3-4 per group) before being immediately sacrificed for tissue collection. The second cohort of rats were used for intra-cortical recordings and perilesional electrophysiological response to the left paw stimulation (see electrophysiological recordings and paw stimulation section; n = 3-4 per group) before being sacrificed for tissue collection. These two cohorts were used to map the volumetric activation of neuronal networks in relation to either volitional control of the left forelimb (SRT cohort) or the sensorimotor response to (Paw Stim). All brains were collected after transcardial perfusion of paraformaldehyde as described above, and processed for iDisco+ tissue clearing.

#### 5. iDisco+ tissue clearing and immunostaining

Following post-fixation, the frontal sections of the brains were treated for immune staining and tissue clearing using the iDisco+ method^63^ with steps described briefly in Table 3.

**Table 3:** iDisco+ tissue clearing steps. For primary and secondary antibody concentration please refer to table 1 and 2. RT: room temperature. DCM: dichloromethane. DBE: Dibenzyl ether. PBS: phosphate buffered saline. PTx2: 0.05% Triton-X100 in 1x PBS. PTwh: 0.0.5% Tween-20, 0.5mL of 10mg/mL of Heparin in 1x PBS.

| Process | Condition | Reagent | Solvent |
| --- | --- | --- | --- |
| Initial wash | 15min x3 in each | Ptx2 and PTwh | 1x PBS |
| Serial dehydration | 1h each | 20/40/60/80/100%<br>methanol | 1x PBS |
| Defatting | Overnight | DCM (66% v/v) | Methanol |
| Bleaching | Overnight | H2O2 (5% v/v) | Methanol |
| Serial rehydration | 1h each | 20/40/60/80/100%<br>PBS | Methanol |
| permeabilization | 4 days | Triton-X100, 20%<br>DMSO | 1x PBS |
| Blocking | 2 days | Goat serum (6%v/v)<br>and DMSO (10% v/v) | Ptx2 |
| Primary antibody<br>incubation | 7 days, 37deg C, 100<br>RPM | Goat serum (3%v/v)<br>and DMSO (5% v/v) | Ptx2 |
| wash | overnight |  | PTwh |
| Secondary antibody<br>incubation | 6 days, 37deg C, 100<br>RPM | Goat serum (3%v/v)<br>and DMSO (5% v/v) | Ptx2 |
| Final dehydration | 1h each | 20/40/60/80/100%<br>methanol | DiH2O |
| defatting | 3h, RT | DCM (66% v/v) | Methanol |
| defatting | Overnight | DCM 100% | DCM |
| Final<br>defatting/clearing | 30min – hold<br>indefinitely | 100% DBE | DBE |

#### 6. Tissue imaging and quantification

Brain slices were imaged using either a Zeiss LSM7100 confocal microscope (Zen^®^ software; Immunostained slices), epifluorescence Leica DM IRBE (Volocity^®^ software; Cresyl/Nissl Stain slices) or Lavision Ultra II lightsheet microscope (iDisco+ cleared brains).

Image quantification for immunostained slices was performed using MATLAB and custom scripts based on the image processing toolbox. For cresyl violet stained slides the region of interest (ROI) represented 10.494 mm^2^, and four images were taken per animal per group (n = 7, 28 total images). ImageJ was used to calculate and analyze the total number of marked nissl bodies, thresholding to signal peak (~200) and using the subtraction tool to eliminate any noise (National Institute of Health, MD). For volumetric images, all image stacks were converted and processed using the software Imaris ^®^ (Bitplane). Nuclear counts (NeuN+ and Arc+ Staining) and colocalization were performed using spot detection and spot colocalization. Vascular density (Reca1+), orientation was performed using a batch sequence of 1) surface mapping, 2) surface model-based masking, 3) masked image Gaussian filter and 4) filament tracing (closed-loop) for vasculature parameter extraction.

### G – Electrophysiological recordings

#### 1. Perilesional electrophysiological recordings

At week 8, rats that received RFA-targeted lesion were induced and maintained under anesthesia using a Ketamine/Xylazine cocktail (100mg/Kg) and placed on a stereotaxic frame. Following sagittal scalp incision, a craniotomy was performed caudally to the injury (AP: 0 mm, ML: 2 to 4mm, relatively to bregma), followed by durotomy to allow for the insertion of a 32 channel silicone probe (Neuronexus). The silicone probe was then inserted in the motor area (position1: CFA, AP: 0 mmm, ML: 2.5 mm) or sensorimotor area (position2: S1, AP: 0 mmm, ML: 3.5 mm) at a depth of 2 mm from the surface of the brain. All recordings were performed with a single shank, 32 channel linear probe with 50 um spacing between recording sites (177 mm^2^) and a maximum span of 1.6 mm (A1×32-6mm-50-177-CM32, 15um thickness, Gen4, lot# P994). The probe reference was used as main reference and a screw positioned above the right cerebellum was used as ground.

Neural data was digitalized and recorded at 20 kHz (unit gain) using a Multichannel systems (MCS) W2100 acquisition module and a wireless headstage (HS32-EXT-0.5mA, 16bit). For spike analysis, the broadband electrophysiological data was real-time bandpass filtered (300-5000 Hz) and a baseline pre-calculated threshold was used as trigger to save spike waveforms and spike event timestamps (threshold: 5 standard deviation). All recordings were performed for 5 min following a 10 min of stabilization.

#### 2. Paw Electrical Stimulation Evoked Neural Activity

Paw electrical stimulation were performed from a separate HS32-EXT-0.5mA headstage to provide isolated stimulation. The stimulating electrode was a stainless-steel needle inserted in the foot pad of the rat paw. Paw stimulations were performed at a fixed intensity (50uA), biphasic electrical pulse (1msec phase 1) optimized to minimize recording artefacts while eliciting triggered neural response. The Paw stimulation pulses were repeated 120 times at a frequency of 1Hz. Each stimulation was monitored using an LED output a synchronized TTL port to the recording system.

### I - Statistical analysis

All statistical analyses were performed using SigmaPlot (SyStat Software, Inc., CA) or Matlab statistical toolbox (Mathworks, inc.). For multiple group comparison (e.g. immunohistochemistry quantifications, Volume or area quantification), we used a one-way ANOVA followed by a multiple comparison least significant difference (LSD) due to the low number of group (3 groups, SHAM, TBI, and eCS). For group comparison on the normalized phase gradient analysis (MRI) we used a 3-way ANOVA followed by multiple comparison using the LSD approach. For behavior data, we performed two-way repeated measure ANOVA using between factor group (three level: SHAM, TBI, eCS) and within factor time with a post-hoc multiple comparison based on the LSD. When necessary (failure of normality of sample distribution), non-parametric tests such as Kruskal-Wallis and Mann-Whitney U Ranksum test were used in place of oneway ANOVA and t-tests. For distribution comparisons, the measure of effect size (Matlab toolbox, dabest estimation statistics)^64^ was used to avoid bias due to high sample number as well as quartile-quartile plot against SHAM distribution. For all tests, *p*<0.05 was considered significant.

## REFERENCES

1 Coronado, V. G., McGuire, L. C., Faul, M., Sugerman, D. & Pearson, W. The epidemiology and prevention of TBI. Brain Injury Medicine. New York, NY: Demos, 45–56 (2012).

2 Pluchino, S. et al. Neurosphere-derived multipotent precursors promote neuroprotection by an immunomodulatory mechanism. Nature 436, 266–271, doi:10.1038/nature03889 (2005).

3 Baker, E. W., Kinder, H. A. & West, F. D. Neural stem cell therapy for stroke: A multimechanistic approach to restoring neurological function. Brain and behavior, e01214 (2019).

4 Wennersten, A., Meijer, X., Holmin, S., Wahlberg, L. & Mathiesen, T. Proliferation, migration, and differentiation of human neural stem/progenitor cells after transplantation into a rat model of traumatic brain injury. Journal of Neurosurgery 100, 88–96, doi:10.3171/jns.2004.100.1.0088 (2004).

5 Liao, G. P. et al. Autologous bone marrow mononuclear cells reduce therapeutic intensity for severe traumatic brain injury in children. Pediatr Crit Care Med 16, 245–255, doi:10.1097/PCC.0000000000000324 (2015).

6 Berman, S. C., Galpoththawela, C., Gilad, A. A., Bulte, J. W. & Walczak, P. Long-term MR cell tracking of neural stem cells grafted in immunocompetent versus immunodeficient mice reveals distinct differences in contrast between live and dead cells. Magn Reson Med 65, 564–574, doi:10.1002/mrm.22613 (2011).

7 Li, J. Y., Christophersen, N. S., Hall, V., Soulet, D. & Brundin, P. Critical issues of clinical human embryonic stem cell therapy for brain repair. Trends Neurosci 31, 146–153, doi:10.1016/j.tins.2007.12.001 (2008).

8 Lee, A. S., Tang, C., Rao, M. S., Weissman, I. L. & Wu, J. C. Tumorigenicity as a clinical hurdle for pluripotent stem cell therapies. Nature medicine 19, 998 (2013).

9 Akita, K. et al. Expression of multiple chondroitin/dermatan sulfotransferases in the neurogenic regions of the embryonic and adult central nervous system implies that complex chondroitin sulfates have a role in neural stem cell maintenance. Stem Cells 26, 798–809, doi:10.1634/stemcells.2007-0448 (2008).

10 Ida, M. et al. Identification and functions of chondroitin sulfate in the milieu of neural stem cells. J Biol Chem 281, 5982–5991, doi:10.1074/jbc.M507130200 (2006).

11 Margolis, R. U., Margolis, R. K., Santella, R. & Atherton, D. M. The hyaluronidase of brain. J Neurochem 19, 2325–2332 (1972).

12 Ruoslahti, E. Structure and biology of proteoglycans. Annu Rev Cell Biol 4, 229–255, doi:10.1146/annurev.cb.04.110188.001305 (1988).

13 Sirko, S., Akita, K., Von Holst, A. & Faissner, A. Structural and functional analysis of chondroitin sulfate proteoglycans in the neural stem cell niche. Methods Enzymol 479, 37–71, doi:10.1016/S0076-6879(10)79003-0 (2010).

14 Sirko, S. et al. Chondroitin sulfates are required for fibroblast growth factor-2-dependent proliferation and maintenance in neural stem cells and for epidermal growth factor-dependent migration of their progeny. Stem Cells 28, 775–787, doi:10.1002/stem.309 (2010).

15 Sirko, S., von Holst, A., Wizenmann, A., Gotz, M. & Faissner, A. Chondroitin sulfate glycosaminoglycans control proliferation, radial glia cell differentiation and neurogenesis in neural stem/progenitor cells. Development 134, 2727–2738, doi:10.1242/dev.02871 (2007).

16 Seo, J. H., Yu, J. H., Suh, H., Kim, M. S. & Cho, S. R. Fibroblast growth factor-2 induced by enriched environment enhances angiogenesis and motor function in chronic hypoxic-ischemic brain injury. PLoS One 8, e74405, doi:10.1371/journal.pone.0074405 (2013).

17 Karumbaiah, L. et al. Chondroitin sulfate glycosaminoglycan hydrogels create endogenous niches for neural stem cells. Bioconjugate chemistry 26, 2336–2349 (2015).

18 Deepa, S. S., Umehara, Y., Higashiyama, S., Itoh, N. & Sugahara, K. Specific molecular interactions of oversulfated chondroitin sulfate E with various heparin-binding growth factors. Implications as a physiological binding partner in the brain and other tissues. J Biol Chem 277, 43707–43716, doi:10.1074/jbc.M207105200 (2002).

19 Nandini, C. D. & Sugahara, K. Role of the sulfation pattern of chondroitin sulfate in its biological activities and in the binding of growth factors. Adv Pharmacol 53, 253–279, doi:10.1016/S1054-3589(05)53012-6 (2006).

20 Carulli, D., Laabs, T., Geller, H. M. & Fawcett, J. W. Chondroitin sulfate proteoglycans in neural development and regeneration. Curr Opin Neurobiol 15, 116–120, doi:10.1016/j.conb.2005.01.014 (2005).

21 Dityatev, A. & Schachner, M. Extracellular matrix molecules and synaptic plasticity. Nat Rev Neurosci 4, 456–468, doi:10.1038/nrn1115 (2003).

22 Frischknecht, R. et al. Brain extracellular matrix affects AMPA receptor lateral mobility and shortterm synaptic plasticity. Nature neuroscience 12, 897 (2009).

23 Gama, C. I. et al. Sulfation patterns of glycosaminoglycans encode molecular recognition and activity. Nat Chem Biol 2, 467–473, doi:10.1038/nchembio810 (2006).

24 Woodbury, M. E. & Ikezu, T. Fibroblast growth factor-2 signaling in neurogenesis and neurodegeneration. J Neuroimmune Pharmacol 9, 92–101, doi:10.1007/s11481-013-9501-5 (2014).

25 Tang, M. M., Lin, W. J., Zhang, J. T., Zhao, Y. W. & Li, Y. C. Exogenous FGF2 reverses depressive-like behaviors and restores the suppressed FGF2-ERK1/2 signaling and the impaired hippocampal neurogenesis induced by neuroinflammation. Brain Behav Immun 66, 322–331, doi:10.1016/j.bbi.2017.05.013 (2017).

26 Dayer, A. G. et al. Expression of FGF-2 in neural progenitor cells enhances their potential for cellular brain repair in the rodent cortex. Brain 130, 2962–2976, doi:10.1093/brain/awm200 (2007).

27 Monfils, M. H. et al. FGF-2-induced cell proliferation stimulates anatomical, neurophysiological and functional recovery from neonatal motor cortex injury. Eur J Neurosci 24, 739–749, doi:10.1111/j.1460-9568.2006.04939.x (2006).

28 Yoshimura, S. et al. FGF-2 regulates neurogenesis and degeneration in the dentate gyrus after traumatic brain injury in mice. J Clin Invest 112, 1202–1210, doi:10.1172/JCI16618 (2003).

29 Cunha, C., Brambilla, R. & Thomas, K. L. A simple role for BDNF in learning and memory? Front Mol Neurosci 3, 1, doi:10.3389/neuro.02.001.2010 (2010).

30 Chen, A., Xiong, L. J., Tong, Y. & Mao, M. The neuroprotective roles of BDNF in hypoxic ischemic brain injury. Biomed Rep 1, 167–176, doi:10.3892/br.2012.48 (2013).

31 Griesbach, G. S., Hovda, D. A., Molteni, R., Wu, A. & Gomez-Pinilla, F. Voluntary exercise following traumatic brain injury: brain-derived neurotrophic factor upregulation and recovery of function. Neuroscience 125, 129–139, doi:10.1016/j.neuroscience.2004.01.030 (2004).

32 Xuan, W., Agrawal, T., Huang, L., Gupta, G. K. & Hamblin, M. R. Low-level laser therapy for traumatic brain injury in mice increases brain derived neurotrophic factor (BDNF) and synaptogenesis. J Biophotonics 8, 502–511, doi:10.1002/jbio.201400069 (2015).

33 Chu, H., Gao, J., Chen, C. W., Huard, J. & Wang, Y. Injectable fibroblast growth factor-2 coacervate for persistent angiogenesis. Proc Natl Acad Sci U S A 108, 13444–13449, doi:10.1073/pnas.1110121108 (2011).

34 Fouda, A. Y. et al. Brain-Derived Neurotrophic Factor Knockdown Blocks the Angiogenic and Protective Effects of Angiotensin Modulation After Experimental Stroke. Mol Neurobiol 54, 661–670, doi:10.1007/s12035-015-9675-3 (2017).

35 Kermani, P. & Hempstead, B. BDNF Actions in the Cardiovascular System: Roles in Development, Adulthood and Response to Injury. Front Physiol 10, 455, doi:10.3389/fphys.2019.00455 (2019).

36 Betancur, M. I. et al. Chondroitin Sulfate Glycosaminoglycan Matrices Promote Neural Stem Cell Maintenance and Neuroprotection Post-Traumatic Brain Injury, 2017).

37 Betancur, M. I. et al. Chondroitin sulfate glycosaminoglycan matrices promote neural stem cell maintenance and neuroprotection post-traumatic brain injury. ACS biomaterials science & engineering 3, 420–430 (2017).

38 Walker, P. A., Shah, S. K., Harting, M. T. & Cox, C. S., Jr. Progenitor cell therapies for traumatic brain injury: barriers and opportunities in translation. Dis Model Mech 2, 23–38, doi:10.1242/dmm.001198 (2009).

39 Nih, L. R., Carmichael, S. T. & Segura, T. Hydrogels for brain repair after stroke: an emerging treatment option. Curr Opin Biotechnol 40, 155–163, doi:10.1016/j.copbio.2016.04.021 (2016).

40 Janz, R. et al. Essential roles in synaptic plasticity for synaptogyrin I and synaptophysin I. Neuron 24, 687–700 (1999).

41 Kumar, A. & Loane, D. J. Neuroinflammation after traumatic brain injury: opportunities for therapeutic intervention. Brain Behav Immun 26, 1191–1201, doi:10.1016/j.bbi.2012.06.008 (2012).

42 Moon, J. J. et al. Biomimetic hydrogels with pro-angiogenic properties. Biomaterials 31, 3840–3847, doi:10.1016/j.biomaterials.2010.01.104 (2010).

43 Goldenberg, G., Oder, W., Spatt, J. & Podreka, I. Cerebral correlates of disturbed executive function and memory in survivors of severe closed head injury: a SPECT study. J Neurol Neurosurg Psychiatry 55, 362–368, doi:10.1136/jnnp.55.5.362 (1992).

44 Ichise, M. et al. Technetium-99m-HMPAO SPECT, CT and MRI in the evaluation of patients with chronic traumatic brain injury: a correlation with neuropsychological performance. J Nucl Med 35, 217–226 (1994).

45 Oder, W. et al. Behavioural and psychosocial sequelae of severe closed head injury and regional cerebral blood flow: a SPECT study. J Neurol Neurosurg Psychiatry 55, 475–480, doi:10.1136/jnnp.55.6.475 (1992).

46 Prayer, L. et al. Cranial MR imaging and cerebral 99mTc HM-PAO-SPECT in patients with subacute or chronic severe closed head injury and normal CT examinations. Acta Radiol 34, 593–599 (1993).

47 Inoue, Y. et al. Changes in cerebral blood flow from the acute to the chronic phase of severe head injury. J Neurotrauma 22, 1411–1418, doi:10.1089/neu.2005.22.1411 (2005).

48 Kim, J. et al. Resting cerebral blood flow alterations in chronic traumatic brain injury: an arterial spin labeling perfusion FMRI study. J Neurotrauma 27, 1399–1411, doi:10.1089/neu.2009.1215 (2010).

49 Ances, B. M., Zarahn, E., Greenberg, J. H. & Detre, J. A. Coupling of neural activation to blood flow in the somatosensory cortex of rats is time-intensity separable, but not linear. J Cereb Blood Flow Metab 20, 921–930, doi:10.1097/00004647-200006000-00004 (2000).

50 Levine, B. et al. The Toronto traumatic brain injury study: injury severity and quantified MRI. Neurology 70, 771–778, doi:10.1212/01.wnl.0000304108.32283.aa (2008).

51 McCrary, M. R. et al. Cortical Transplantation of Brain-Mimetic Glycosaminoglycan Scaffolds and Neural Progenitor Cells Promotes Vascular Regeneration and Functional Recovery after Ischemic Stroke in Mice. Adv Healthc Mater 9, e1900285, doi:10.1002/adhm.201900285 (2020).

52 Nih, L. R., Gojgini, S., Carmichael, S. T. & Segura, T. Dual-function injectable angiogenic biomaterial for the repair of brain tissue following stroke. Nature Materials 17, 642, doi:10.1038/s41563-018-0083-8 (2018).

53 Nudo, R. J. Adaptive plasticity in motor cortex: implications for rehabilitation after brain injury. J Rehabil Med, 7–10 (2003).

54 Nudo, R. J. Mechanisms for recovery of motor function following cortical damage. Current Opinion in Neurobiology 16, 638–644, doi:10.1016/j.conb.2006.10.004 (2006).

55 Kim, S. Y., Hsu, J. E., Husbands, L. C., Kleim, J. A. & Jones, T. A. Coordinated Plasticity of Synapses and Astrocytes Underlies Practice-Driven Functional Vicariation in Peri-Infarct Motor Cortex. Journal of Neuroscience 38, 93–107, doi:10.1523/JNEUROSCI.1295-17.2017 (2018).

56 Ramanathan, D. S. et al. Low-frequency cortical activity is a neuromodulatory target that tracks recovery after stroke. Nature Medicine 24, 1257, doi:10.1038/s41591-018-0058-y (2018).

57 Fujimoto, S. T. et al. Motor and cognitive function evaluation following experimental traumatic brain injury. Neurosci Biobehav Rev 28, 365–378, doi:10.1016/j.neubiorev.2004.06.002 (2004).

58 Hamm, R. J., Pike, B. R., O’Dell, D. M., Lyeth, B. G. & Jenkins, L. W. The rotarod test: an evaluation of its effectiveness in assessing motor deficits following traumatic brain injury. J Neurotrauma 11, 187–196, doi:10.1089/neu.1994.11.187 (1994).

59 Shen, Y. et al. In vivo measurement of tissue damage, oxygen saturation changes and blood flow changes after experimental traumatic brain injury in rats using susceptibility weighted imaging. Magnetic resonance imaging 25, 219–227 (2007).

60 Wang, L., Potter, W. M. & Zhao, Q. In vivo quantification of SPIO nanoparticles for cell labeling based on MR phase gradient images. Contrast media & molecular imaging 10, 43–50 (2015).

61 Simchick, G., Betancur, M., Karumbaiah, L. & Zhao, Q. Gauging the Effectiveness of Traumatic Brain Injury Treatment using MR Phase Gradient Mapping. In Proceedings of the 24th Annual Meeting of ISMRM, Singapore, Singapore (2016).

62 Karumbaiah, L. et al. The upregulation of specific interleukin (IL) receptor antagonists and paradoxical enhancement of neuronal apoptosis due to electrode induced strain and brain micromotion. Biomaterials 33, 5983–5996, doi:10.1016/j.biomaterials.2012.05.021 (2012).

63 Renier, N. et al. iDISCO: a simple, rapid method to immunolabel large tissue samples for volume imaging. Cell 159, 896–910, doi:10.1016/j.cell.2014.10.010 (2014).

64 Ho, J., Tumkaya, T., Aryal, S., Choi, H. & Claridge-Chang, A. Moving beyond P values: data analysis with estimation graphics. Nature Methods 16, 565–566 (2019).

