## Supplemental Figures and Videos for "Neurotrophic Factor-Laden Acellular Chondroitin Sulfate Scaffolds Promote Chronic Functional Recovery After Severe Traumatic Brain Injury"

**Correspondence:**

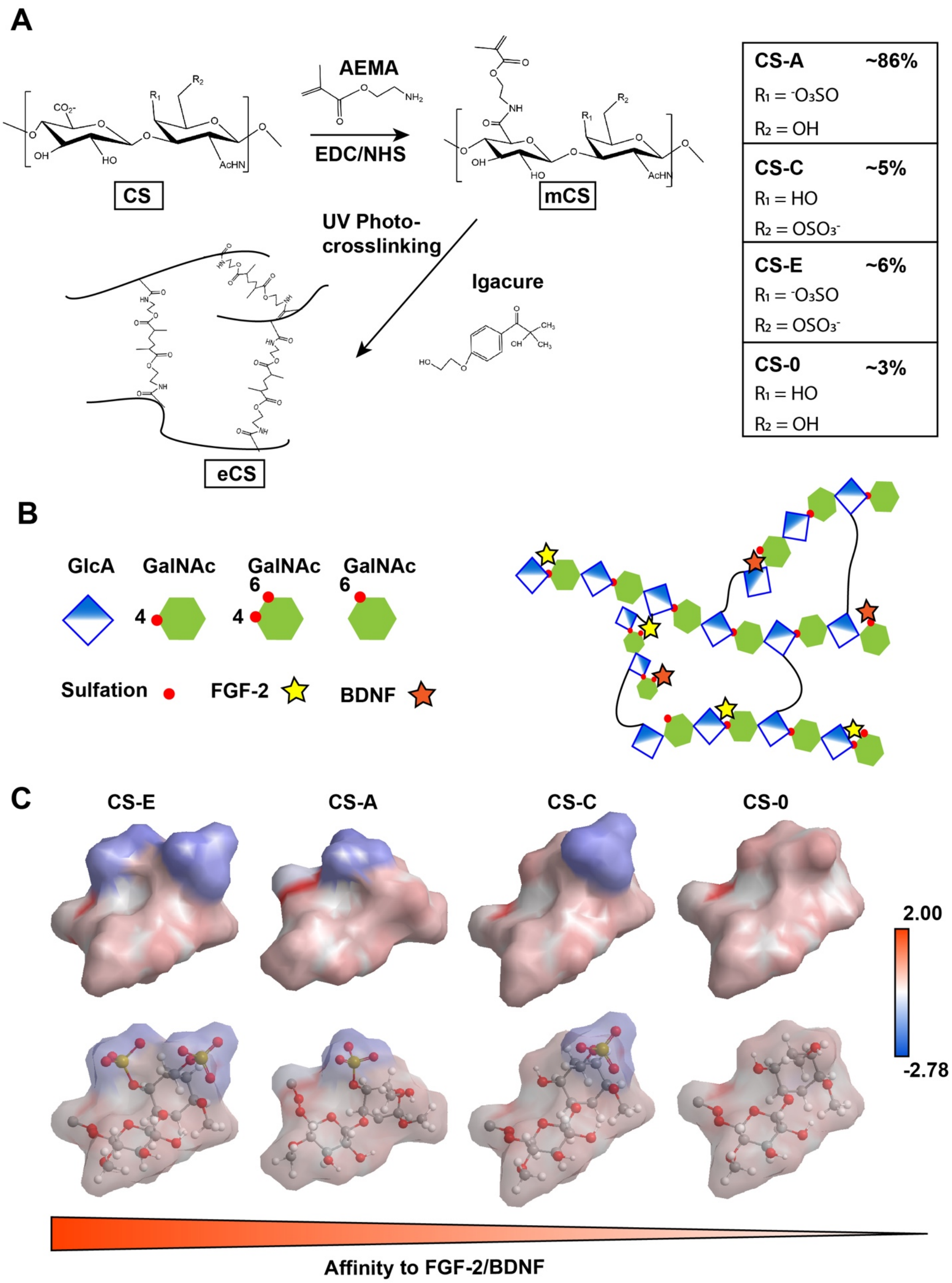

A
AEMA
EDC/NHS
CS
mCS
UV Photo-crosslinking
Igacure
eCS
CS-A ~86%
R1 = -O3SO
R2 = OH
CS-C ~5%
R1 = HO
R2 = OSO3-
CS-E ~6%
R1 = -O3SO
R2 = OSO3-
CS-0 ~3%
R1 = HO
R2 = OH
B
GlcA
GalNAc
Sulfation
FGF-2
BDNF
C
CS-E
CS-A
CS-C
CS-0
2.00
-2.78
Affinity to FGF-2/BDNF

**Supplementary Figure 1** – eCS hydrogel synthesis and sulfation dependent affinity binding of FGF-2 and BDNF

A – Polymeric CS consisting of different percentages of monosulfated (4-O sulfate; CS-A, and 6-O sulfate; CS-C), di-sulfated (4,6-O-culfate; CS-E), and unsulfated (CS-0) CS disaccharides (presented in inset table, right panel), was carboxyl ion functionalized with 2-aminoethyl methacrylate (AEMA) using carbodiimide chemistry. The methacrylate derivatized CS (mCS; 5% w/v) was mixed in PBS containing FGF-2 (55ng) and BDNF (55 ng) along with photocrosslinker (0.1% w/v Irgacure 2959), and exposed to long-wave UV (365nm) to form hydrogels.

B – Schematic representation of FGF-2 and BDNF bound to mono- and disulfated CS motifs in the meshwork of eCS scaffolds.

C – Hückel calculation (surface type: Connolly) of electron density of CS disaccharides. Atomic charges are represented using graded color from blue (negative) to red (positive). The color-coded surfaces are presented with solid color (top panel) and translucent color (bottom panel) to improve visibility of charge distribution and charged group ball-and-stick representations, respectively.

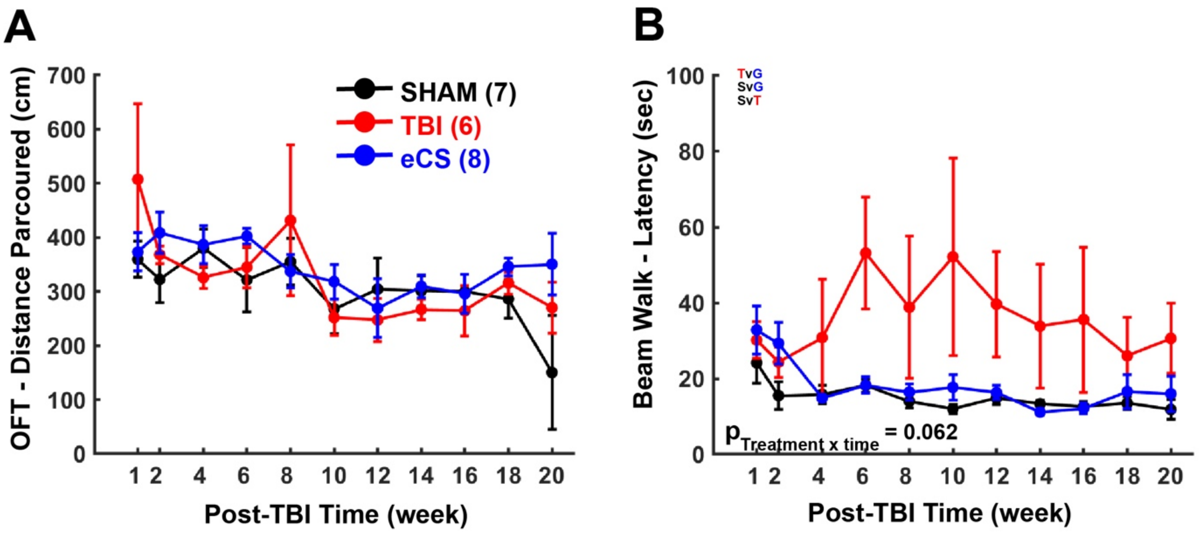


**Supplementary Figure 2** – Hindlimb motor recovery post-TBI targeting CFA (M1) in SHAM, TBI and eCS rats:

A – The open field task (OFT) was performed to assess the basic change in free exploration and locomotion in an open space with higher distance travelled indicating higher locomotion activity. Two-way repeated measure ANOVA; $p_{Treatment} = 0.463$; $p_{Time} < 0.001$; $p_{TreatmentxTime} = 0.010$. Graph show mean ± s.e.m.

B – The beam walk (BW) assay was performed to assess the forced balance on a narrow beam and motor coordination with shorter latency to cross the beam indicating better balance and coordination. Two-way repeated measure ANOVA; $p_{Treatment} = 0.062$; $p_{Time} = 0.069$; $p_{TreatmentxTime} = 0.062$. Graph show mean ± s.e.m.

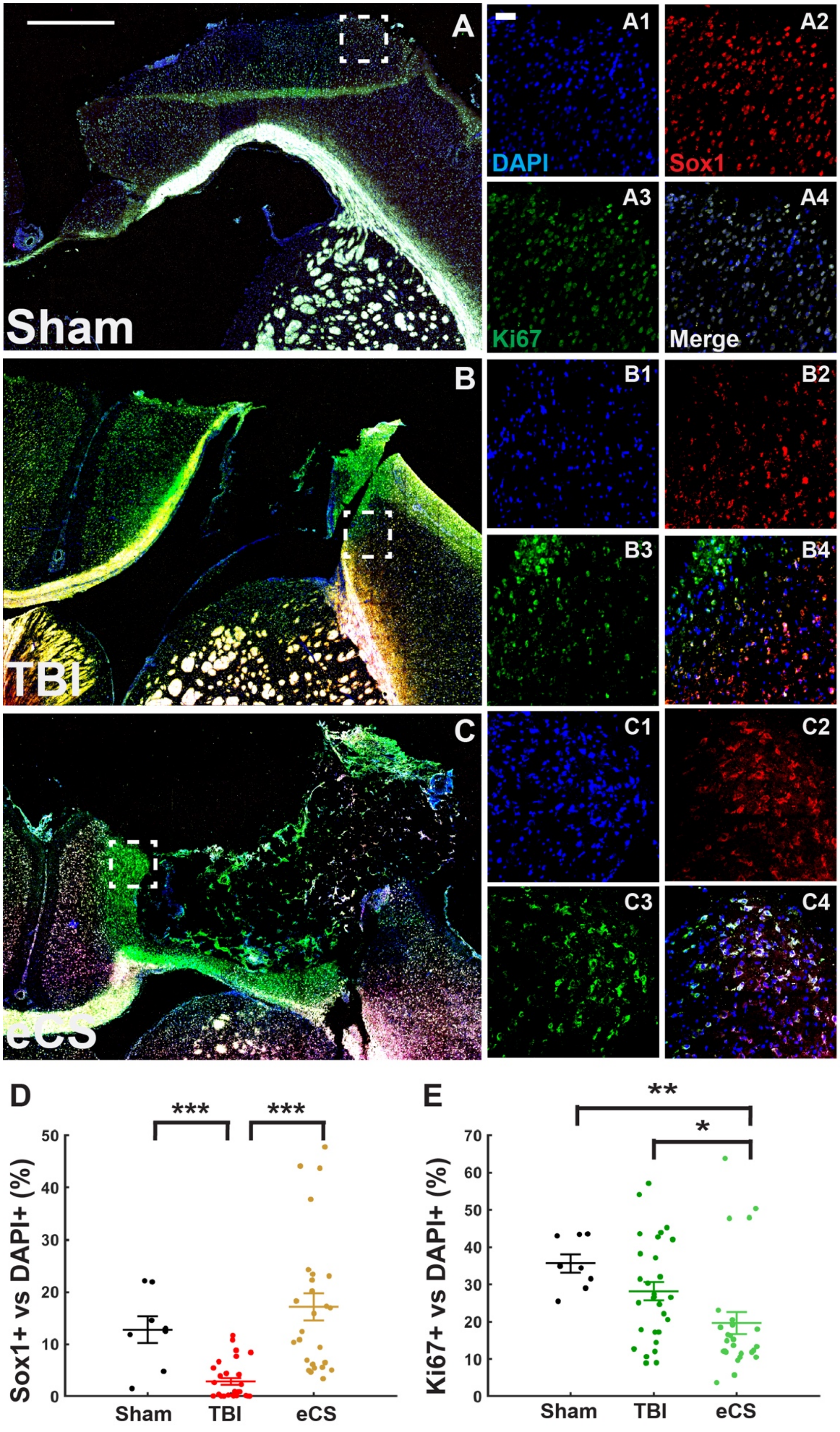
A
A1
A2
DAPI
Sox1
A3
A4
Ki67
Merge
Sham
B
B1
B2
B3
B4
TBI
C
C1
C2
C3
C4
eCS
D
***
***
Sox1+ vs DAPI+ (%)
50
40
30
20
10
0
Sham
TBI
eCS
E
**
*
Ki67+ vs DAPI+ (%)
70
60
50
40
30
20
10
0
Sham
TBI
eCS

**Supplementary Figure 3** – Neural progenitor presence

A-C – Representative tiled images of ipsilesional hemisphere (left coronal sections) for the treatment groups SHAM (top), TBI (middle), eCS (bottom); scale bar = 1mm. A1-A4 – representative magnification of dashed white square shown in A for DAPI (A1), Sox1 (A2), Ki67(A3) and merged (A4); scale bar is 100 µm; B1-B4 – representative magnification of dashed white square shown in A for DAPI (B1), Sox1 (B2), Ki67(B3) and merged (B4); scale bar is 100 µm; C1-C4 – representative magnification of dashed white square shown in C for DAPI (C1), Sox1 (C2), Ki67(C3) and merged (C4); scale bar is 100 µm;

D – Co-localization of Sox1+ cells with DAPI+ cells as percentage of DAPI+ cells for each treatment. Kruskal-Wallis, Treatment: $p < 0.001$.

E – Co-localization of Ki67+ cells with DAPI+ cells as percentage of DAPI+ cells for each treatment. One-way ANOVA, Treatment: $p = 0.0084$. Post-hoc LSD Mann-Whitney U test *, **, and *** are for $p<0.05$, $p<0.01$, and $p<0.001$, respectively. Graphs show mean ± s.e.m.

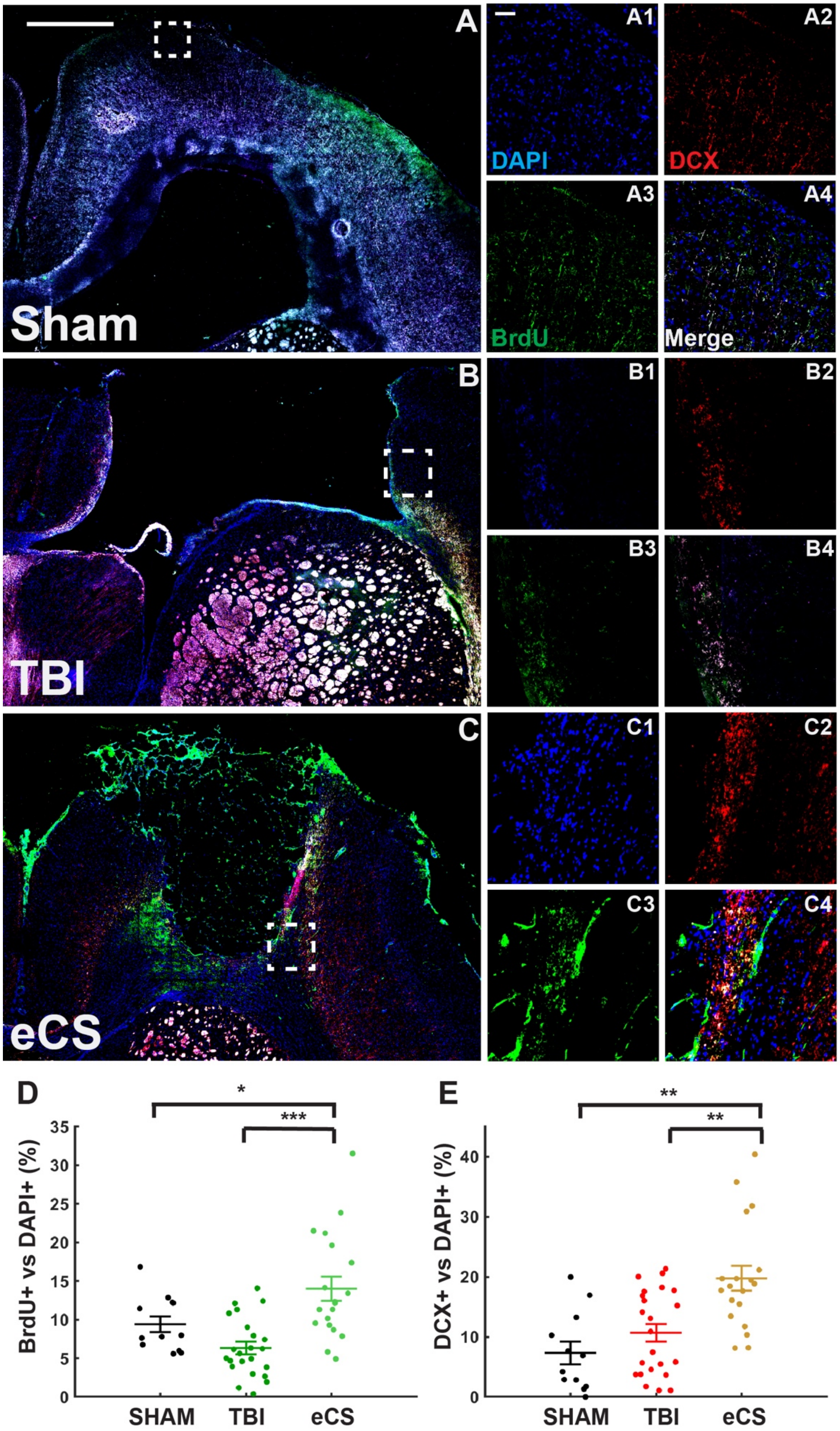
A
Sham
A1
A2
A3
A4
DAPI
DCX
BrdU
Merge
B
TBI
B1
B2
B3
B4
C
eCS
C1
C2
C3
C4
D
*
***
BrdU+ vs DAPI+ (%)
SHAM
TBI
eCS
E
**
**
DCX+ vs DAPI+ (%)
SHAM
TBI
eCS

**Supplementary Figure 4** – Neurogenesis Markers

A-C – Representative tiled images of ipsilesional hemisphere (left coronal sections) sections for the treatment groups SHAM (top), TBI (middle), eCS (bottom); scale bar = 1mm.

A1-A4 – representative magnification of dashed white square shown in A for DAPI (A1), DCX (A2), BrdU(A3) and merged (A4); scale bar is 100 µm;

B1-B4 – representative magnification of dashed white square shown in A for DAPI (B1), DCX (B2), BrdU(B3) and merged (B4); scale bar is 100 µm;

C1-C4 – representative magnification of dashed white square shown in C for DAPI (C1), DCX (C2), BrdU(C3) and merged (C4); scale bar is 100 µm;

D – Co-localization of BrdU+ cells with DAPI+ cells as percentage of DAPI+ cells for each treatment. One-way ANOVA, Treatment: $p < 0.001$.

E – Co-localization of DCX+ cells with DAPI+ cells as percentage of DAPI+ cells for each treatment; Kruskal-Wallis, Treatment: $p < 0.001$. Post-hoc LSD Mann-Whitney U test *, **, and *** are for $p<0.05$, $p<0.01$, and $p<0.001$, respectively. Graphs show mean ± s.e.m.

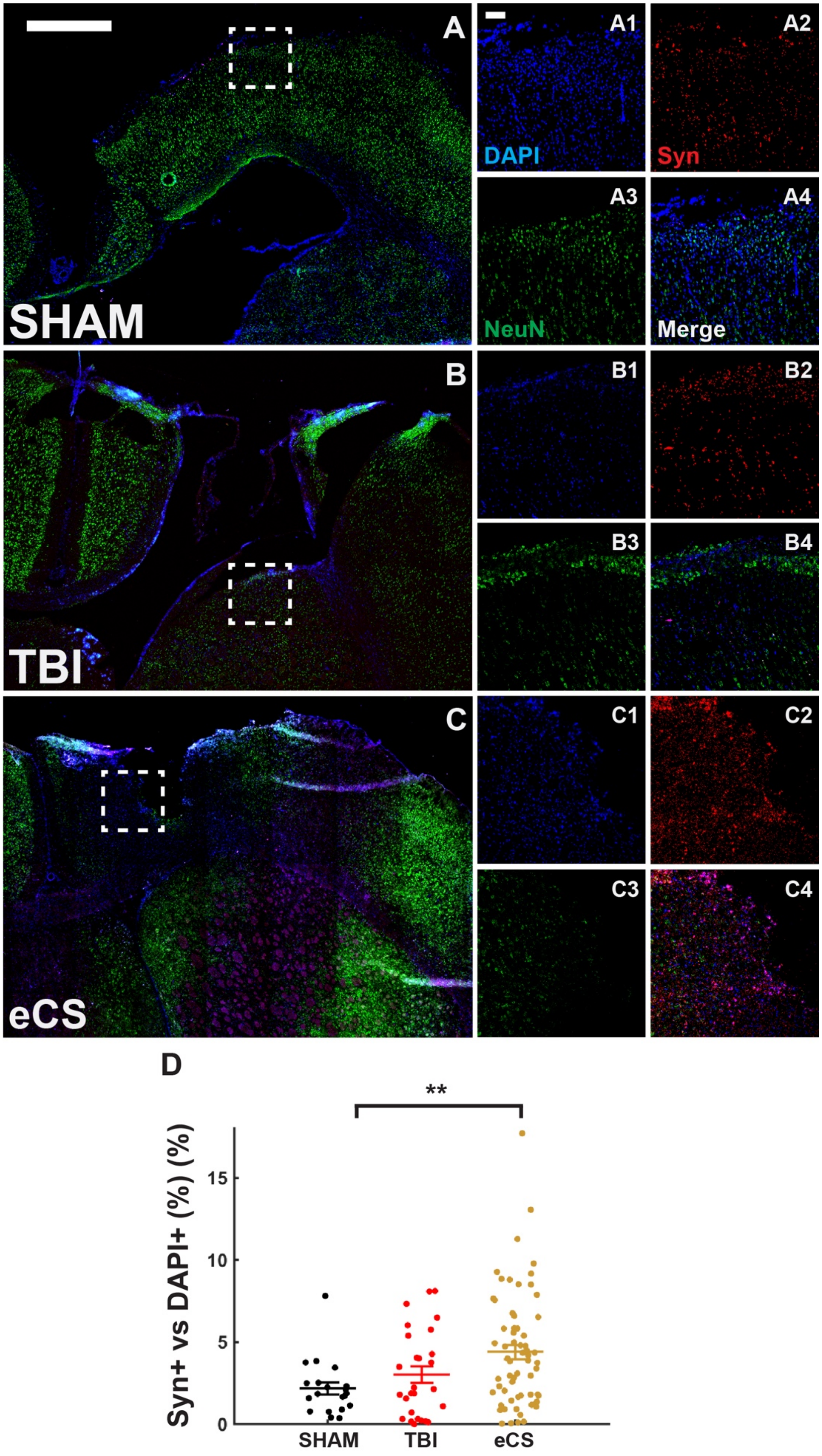

A
A1
A2
DAPI
Syn
A3
A4
NeuN
Merge
SHAM
B
B1
B2
B3
B4
TBI
C
C1
C2
C3
C4
eCS
D
**
Syn+ vs DAPI+ (%) (%)
15
10
5
0
SHAM
TBI
eCS

**Supplementary Figure 5** – Synaptogenesis marker

A-C – Representative tiled images of ipsilesional hemisphere (left coronal sections) for the treatment groups SHAM (top), TBI (middle), eCS (bottom); scale bar = 1mm.

A1-A4 – representative magnification of dashed white square shown in A for DAPI (A1), Syn (A2), NeuN (A3) and merged (A4); scale bar is 100 µm;

B1-B4 – representative magnification of dashed white square shown in A for DAPI (B1), Syn (B2), NeuN (B3) and merged (B4); scale bar is 100 µm;

C1-C4 – representative magnification of dashed white square shown in C for DAPI (C1), Syn (C2), NeuN (C3) and merged (C4); scale bar is 100 µm;

D – Co-localization of Syn + cells with DAPI+ cells as percentage of DAPI+ cells for each treatment. Kruskal-Wallis, Treatment: $p = 0.014$. Post-hoc LSD Mann-Whitney U test *, **, and *** are for $p<0.05$, $p<0.01$, and $p<0.001$, respectively. Graph shows mean ± s.e.m.

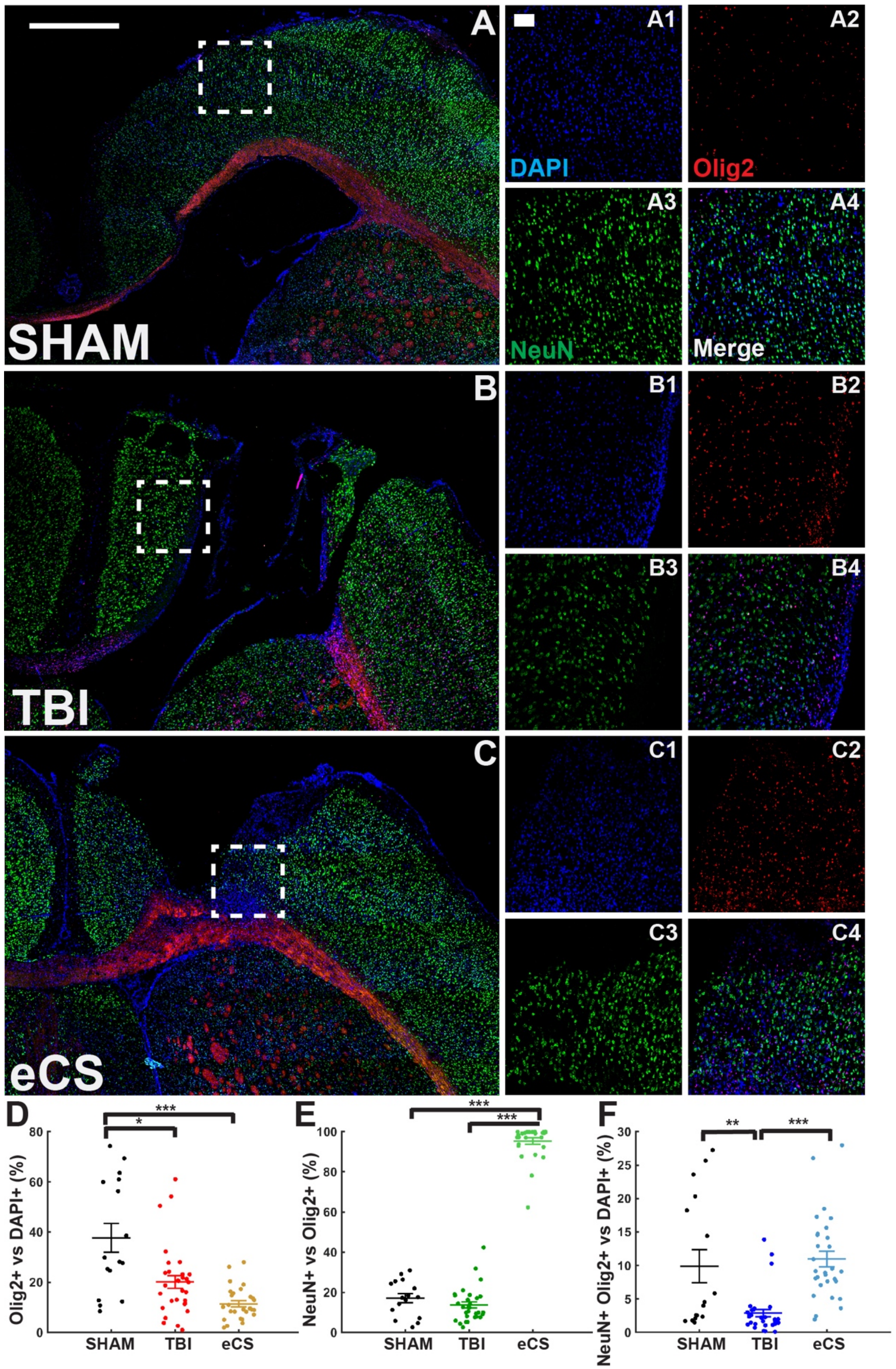

A
A1
A2
DAPI
Olig2
A3
A4
NeuN
Merge
SHAM
B
B1
B2
B3
B4
TBI
C
C1
C2
C3
C4
eCS
D
Olig2+ vs DAPI+ (%)
SHAM
TBI
eCS
E
NeuN+ vs Olig2+ (%)
SHAM
TBI
eCS
F
NeuN+ Olig2+ vs DAPI+ (%)
SHAM
TBI
eCS

**Supplementary Figure 6** – Oligodendrocyte and Neuronal presence

A-C – Representative tiled images of ipsilesional hemisphere (left coronal sections) for the treatment groups SHAM (top), TBI (middle), eCS (bottom); scale bar = 1mm. A1-A4 – representative magnification of dashed white square shown in A for DAPI (A1), Olig2 (A2), NeuN(A3) and merged (A4); scale bar is 100 µm; B1-B4 – representative magnification of dashed white square shown in A for DAPI (B1), Olig2 (B2), NeuN (B3) and merged (B4); scale bar is 100 µm; C1-C4 – representative magnification of dashed white square shown in C for DAPI (C1), Olig2 (C2), NeuN (C3) and merged (C4); scale bar is 100 µm;

D – Co-localization of Olig2+ cells with DAPI+ cells as percentage of DAPI+ cells for each treatment. Kruska-Wallis, Treatment: $p < 0.001$.

E – Co-localization of NeuN + cells with DAPI+ cells as percentage of DAPI+ cells for each treatment. Kruska-Wallis, Treatment: $p < 0.001$.

F – Co-localization of Olig2+ cells with NeuN + cells as percentage of NeuN + cells for each treatment. Kruska-Wallis, Treatment: $p < 0.001$. Post-hoc LSD Mann-Whitney U test *, **, and *** are for $p<0.05$, $p<0.01$, and $p<0.001$, respectively. Graphs show mean ± s.e.m.

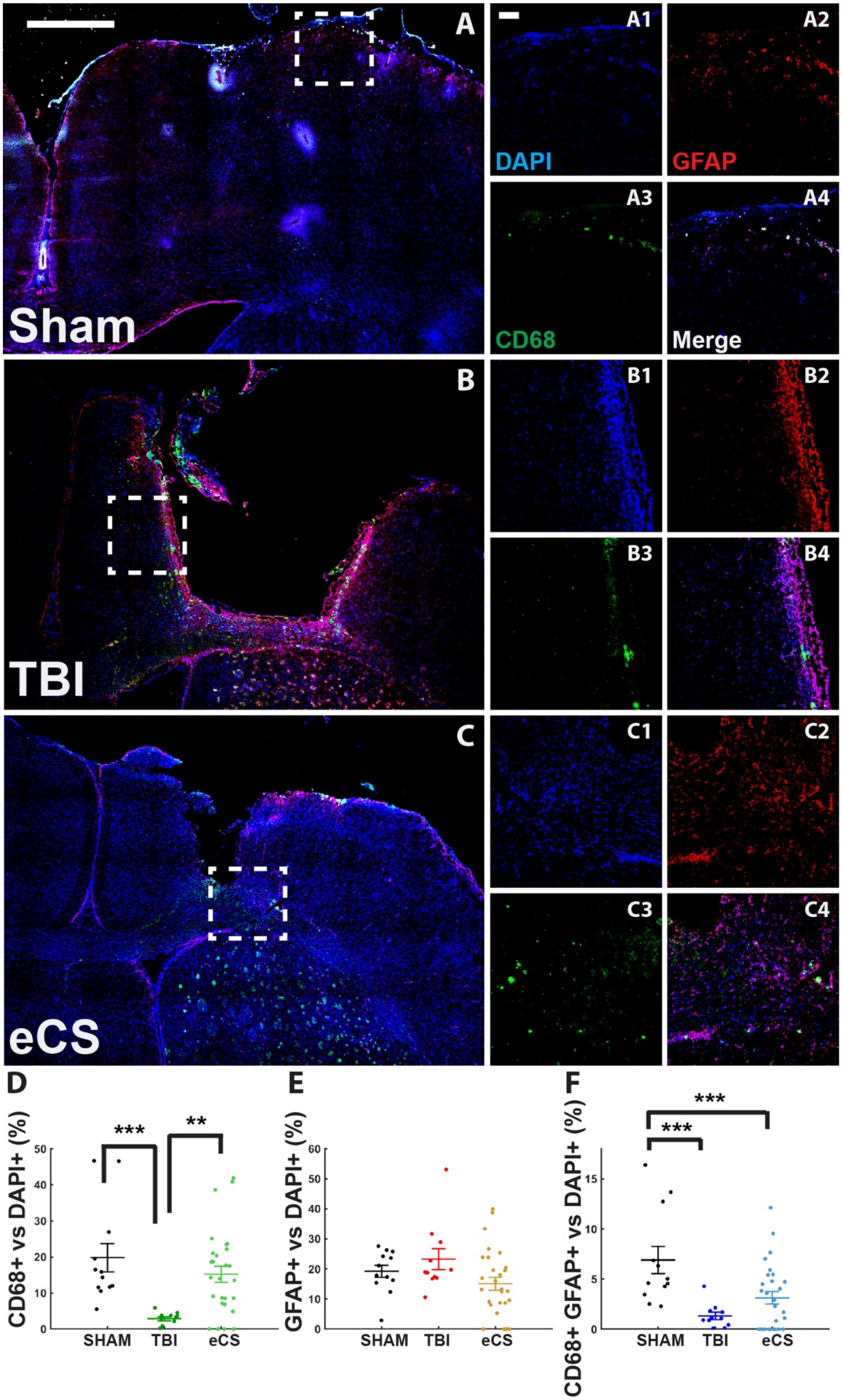
A
A1
A2
A3
A4
DAPI
GFAP
CD68
Merge
Sham
B
B1
B2
B3
B4
TBI
C
C1
C2
C3
C4
eCS
D
E
F
CD68+ vs DAPI+ (%)
GFAP+ vs DAPI+ (%)
CD68+ GFAP+ vs DAPI+ (%)
***
**
***
***
SHAM
TBI
eCS

**Supplementary Figure 7** – eCS promotes the return of neuroimmune response to Sham level

A-C – Representative tiled images of ipsilesional hemisphere (left coronal sections) for the treatment groups Sham (top), TBI (middle), eCS (bottom); scale bar = 1mm. A1-A4 – representative magnification of dashed white square shown in A for DAPI (A1), GFAP (A2), CD68 (A3) and merged (A4); scale bar is 100 µm. B1-B4 – representative magnification of dashed white square shown in A for DAPI (B1), GFAP (B2), CD68 (B3) and merged (B4); scale bar is 100 µm. C1-C4 – representative magnification of dashed white square shown in C for DAPI (C1), GFAP (C2), CD68 (C3) and merged (C4); scale bar is 100 µm;

D – Co-localization CD68+ cells with DAPI+ cells as percentage of DAPI+ cells for each treatment. Kruskal-Wallis, Treatment: $p < 0.001$.

E – Co-localization GFAP+ cells with DAPI+ cells as percentage of DAPI+ cells for each treatment. Kruskal-Wallis, Treatment: $p = 0.094$.

F – Co-localization CD68+ cells with GFAP+ cells as percentage of DAPI+ cells for each treatment. Kruskal-Wallis, Treatment: $p = 0.0033$. Post-hoc LSD Mann-Whitney U test *, **, and *** are for $p<0.05$, $p<0.01$, and $p<0.001$, respectively. Graphs show mean ± s.e.m.

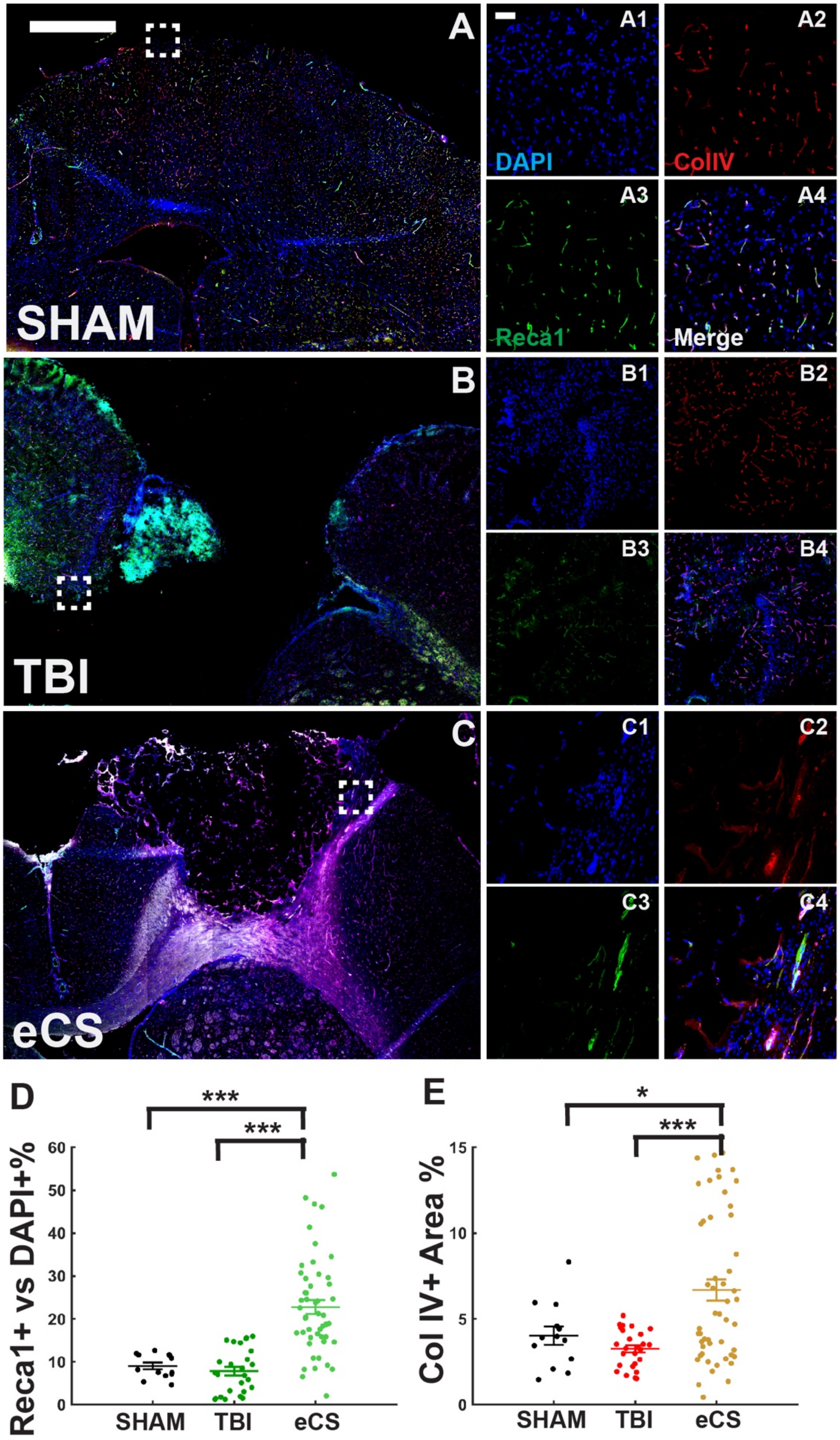
A
A1
A2
DAPI
CollV
A3
A4
Reca1
Merge
SHAM
B
B1
B2
B3
B4
TBI
C
C1
C2
C3
C4
eCS
D
***
***
Reca1+ vs DAPI+%
60
50
40
30
20
10
0
SHAM
TBI
eCS
E
*
***
Col IV+ Area %
15
10
5
0
SHAM
TBI
eCS

**Supplementary Figure 8** – eCS promote long-term Vascularization 20 weeks post-TBI

A-C – Representative tiled images of ipsilesional hemisphere (left coronal sections) for the treatment groups SHAM (top), TBI (middle), eCS (bottom); scale bar = 1mm. A1-A4 – representative magnification of dashed white square shown in A for DAPI (A1), Col IV (A2), Reca1 (A3) and merged (A4); scale bar is 100 µm; B1-B4 – representative magnification of dashed white square shown in A for DAPI (B1), Col IV (B2), Reca1 (B3) and merged (B4); scale bar is 100 µm; C1-C4 – representative magnification of dashed white square shown in C for DAPI (C1), Col IV (C2), Reca1 (C3) and merged (C4); scale bar is 100 µm;

D – Co-localization of Reca1 + cells with DAPI+ cells as percentage of DAPI+ cells for each treatment. One-Way ANOVA, Treatment: $p < 0.001$.

E – Col IV+ are represented as a percentage of image area. One-Way ANOVA, Treatment: $p < 0.001$. Post-hoc LSD Mann-Whitney U test *, **, and *** are for $p<0.05$, $p<0.01$, and $p<0.001$, respectively. Graphs show mean ± s.e.m.

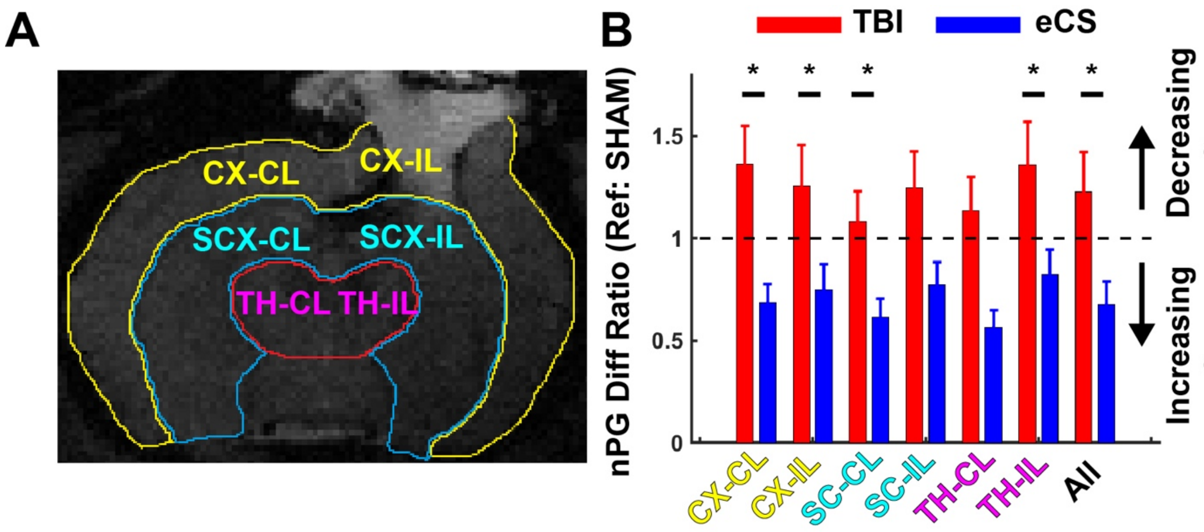


**Supplementary Figure 9** – MRI-based rCBF with subregional quantification

A – Region of Interest defined based on T2W1 maps and used to locate all blood vessels and nPG calculation for each groups.

B – nPG for the TBI and eCS groups normalized to the SHAM. IL: ipsilesional, CL: contralesional, CX: cortex, SCX: sub-cortex/midbrain, TH: thalamus. Post-hoc LSD *, ** and *** indicate a $p<0.05$, $p<0.01$ and $p <0.001$, respectively. Bar graphs show mean ± s.e.m.

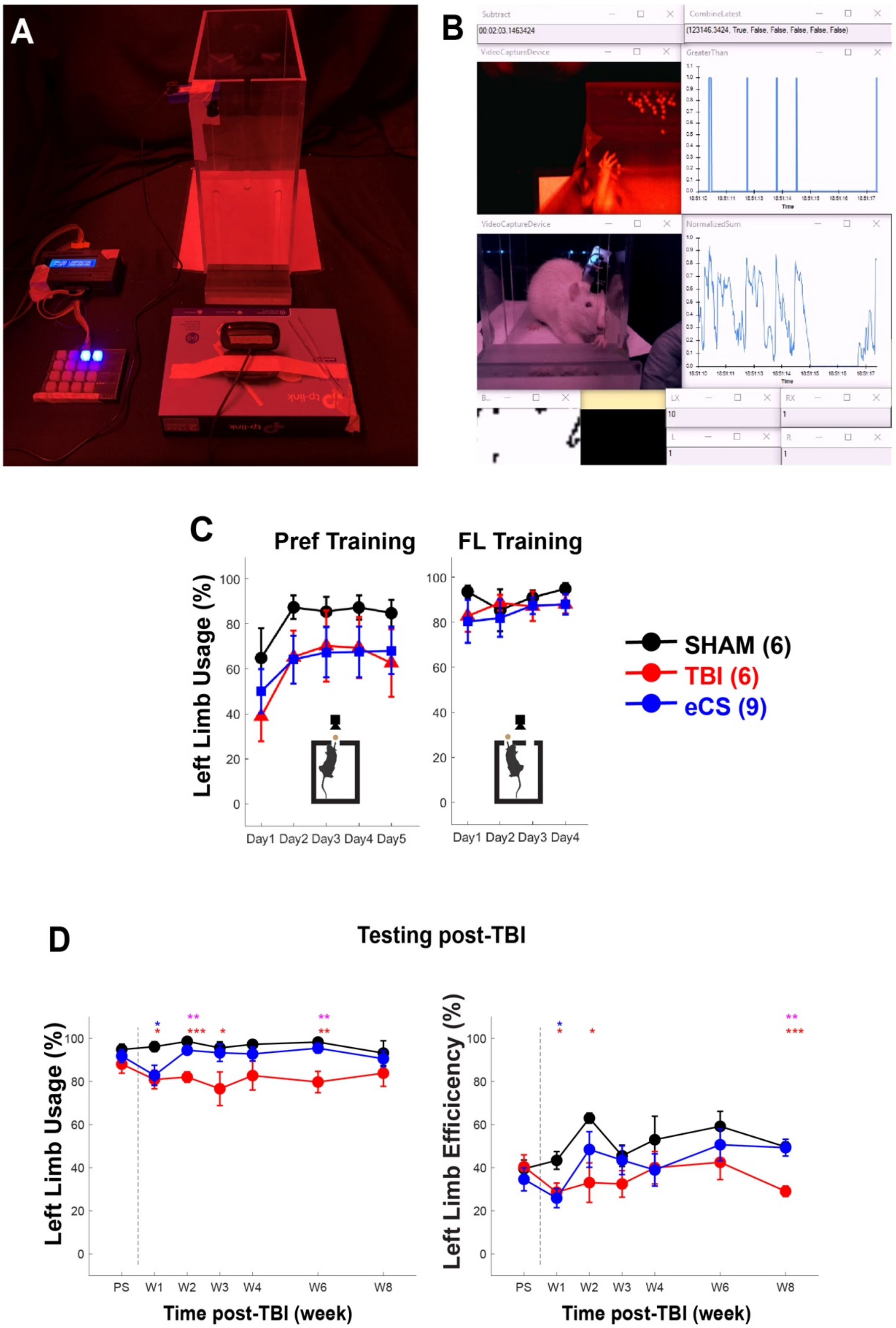
A
B
Subtract
CombineLatest
VideoCaptureDevice
GreaterThan
NormalizedSum
LX
RX
L
R
C
Pref Training
FL Training
Left Limb Usage (%)
Day1 Day2 Day3 Day4 Day5
Day1 Day2 Day3 Day4
SHAM (6)
TBI (6)
eCS (9)
D
Testing post-TBI
Left Limb Usage (%)
Left Limb Efficiency (%)
PS W1 W2 W3 W4 W6 W8
Time post-TBI (week)

**Supplementary Figure 10** – Skilled reach task setup and separate index of performance

A – Picture of the skilled reach task setup. A main Plexiglass box was used for the training and test for preference (one opening side) and forced left (2 opening sides). A custom-made scoring pad was used to score all trials for success, failure and tongue usage. All recordings were sync with a computer, a front and top camera to capture hand movements

B – Snapshot of the recording system using the open source software Bonsai. An Arduino interface was receiving the manual scoring commands while and ROI on the top camera was used to detect individual grabbing tentative from the rat. All data were saved in both video and text file for later analysis.

C - Following preference and forced left limb training, all the rats acquired the usage of their left limb to retrieve the food pellet. Group difference at the end of the preference test indicates the presence of right handed rats. FL: Forced Left assay.

D – Separate score for the left-limb usage (%) and efficiency (%). Post-hoc LSD *, ** and *** indicate a $p<0.05$, $p<0.01$ and $p <0.001$, respectively. The color of stars indicates statistical tests between the following pairs: SHAM vs TBI (SvT, red), SHAM vs eCS (SvG, Blue) and TBI vs eCS (TvG, magenta). Line plots and graphs show mean ± s.e.m.

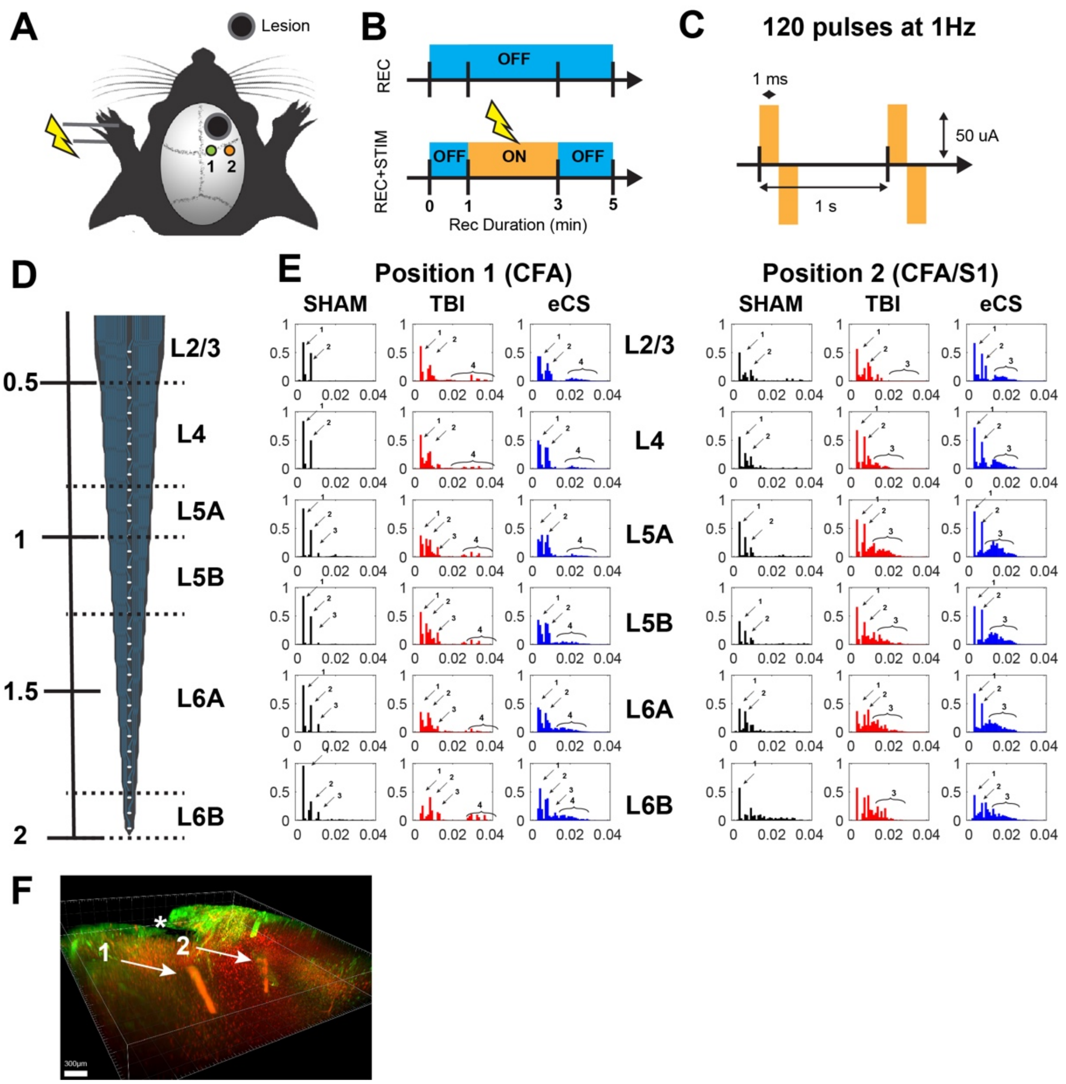
A
Lesion
1 2
B
REC
OFF
REC+STIM
OFF ON OFF
0 1 3 5
Rec Duration (min)
C
120 pulses at 1Hz
1 ms
50 uA
1 s
D
L2/3
0.5
L4
L5A
1
L5B
1.5
L6A
L6B
2
E
Position 1 (CFA)
Position 2 (CFA/S1)
SHAM TBI eCS
L2/3
L4
L5A
L5B
L6A
L6B
F
300μm

**Supplementary Figure 11** – eCS implants promote sensorimotor connectivity 10 weeks post-TBI

A – Experimental schedule for recording session in ketamine anesthetized rats. Two positions for recording were used perilesionally for each rat in order to avoid the disruption of the injury or eCS implant. Position 1: CFA; Position 2: CFA/S1.

B – Recording session for each position was performed first recording only after the probe insertion has been stabilized (top) then a stimulation session (2 min) with pre and post stimulation baseline recording of 1 and 2min, respectively. 120 stimulation pulses were delivered for each position and each rat.

C – Stimulation protocol used a bipolar pulse of 1ms width (phase1) at 1Hz with an amplitude fix for all rats at 50uA. The stimulation amplitude was determined to minimize stimulation artifacts on recording while eliciting sensorimotor response. Note: stimulations and recordings were performed using two separate wireless headstages (Multichannel Systems, W2100 HS32-Ext0.5mA) to guarantee ground and stimulation isolation from recording electrodes. Each rats had recording session without (5 min) and with stimulation (5min, out of which 2min where stimulated) for the two positions CSA and CSA/S1.

D – A 32 channel linear silicone electrode was used (iridium-iridium oxide, recording sites: 50 µm spacing) with a total span of ~1.6 mm. Implantation was performed up to a depth of 2mm from the surface of the pia. The layer position was based on the depth of the electrode and previously characterized layer distribution in the sensorimotor cortex.

E – For the two recording positions, we observed a treatment- and layer-dependent stimulation-locked response. Typically, following stimulation, a multimodal distribution revealed two major sharp peaks of neuronal activity (mono/di-synaptic; arrow indicating peak 1 and 2) and later response that reveals multi-synaptic activation of the area post-Paw stimulation (arrow indicating peak 3 and 4). Arrows indicate detected peaks of activity response, numbered in order of delay from stimulation start.

F – Representative localization of the electrode positioning in a TBI rat brain, post-recording/stimulation. The arrow indicates the position of recording for position 1 and 2 shown in A. * indicates the position of the lesion. Scale bar: 300 µm

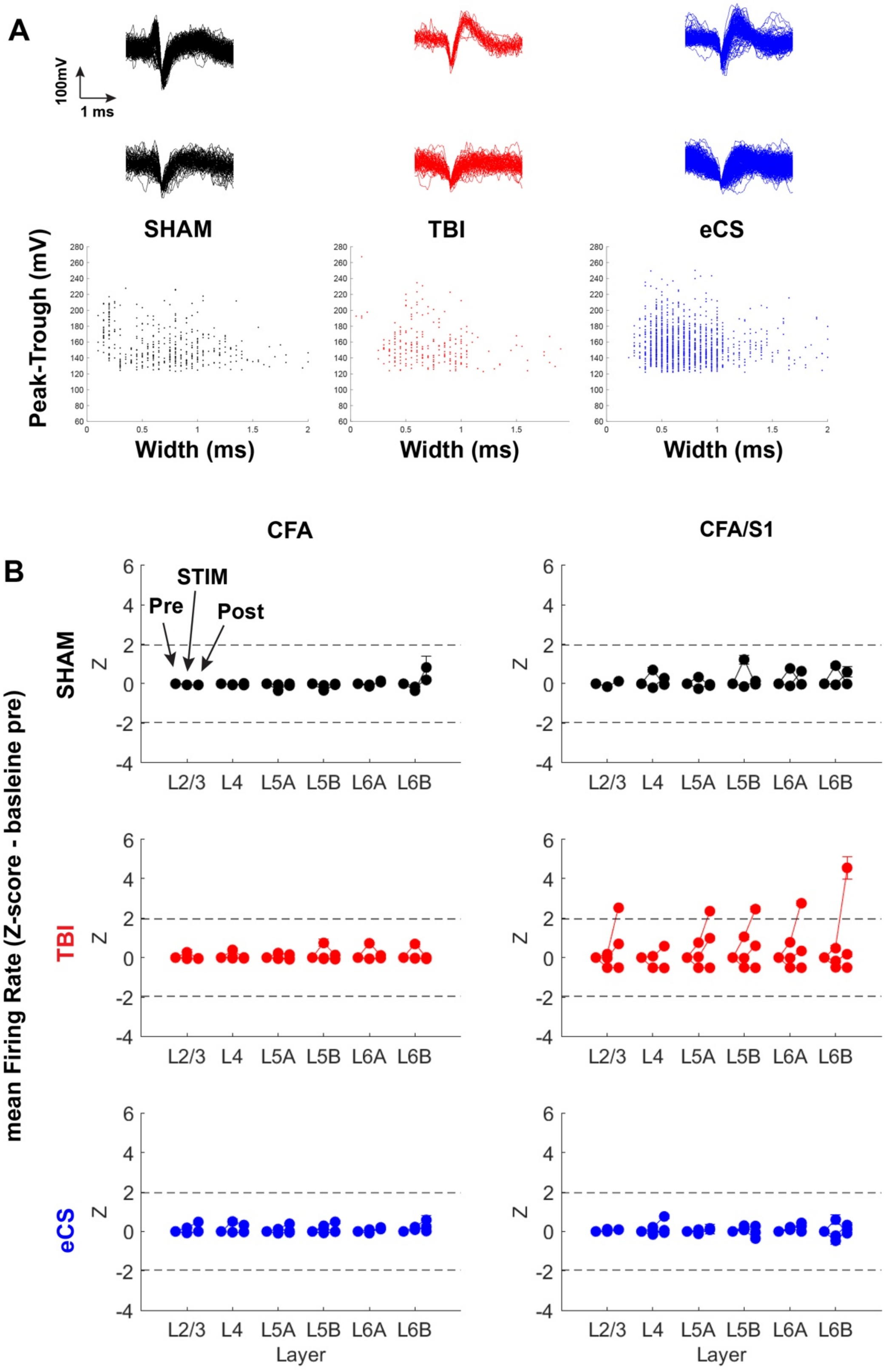
A
100mV
1 ms
SHAM
TBI
eCS
Peak-Trough (mV)
Width (ms)
B
CFA
CFA/S1
STIM
Pre
Post
SHAM
TBI
eCS
mean Firing Rate (Z-score - basleine pre)
Z
L2/3 L4 L5A L5B L6A L6B
Layer

**Supplementary Figure 12** – Characterization of multi-unit activity in all treatment groups using linear probe recording under Ketamine.

A – Multi-unit spike wave form extracted during the resting period for each treatment group for all recording sites. Top panel shows two representative waveforms for each treatment group. Bottom panel shows the width vs peak-trough length scatter plot for all detected multi-unit.

B – Z-score derived from the average population firing frequency normalized to the pre-stimulation period (1min) for each treatment and the two recording positions CFA and CFA/S1/ CFA: caudal forelimb area. Note: TBI group showed a maladaptive sustained firing post stimulation for two rats out of three. Graphs show mean ± s.e.m.

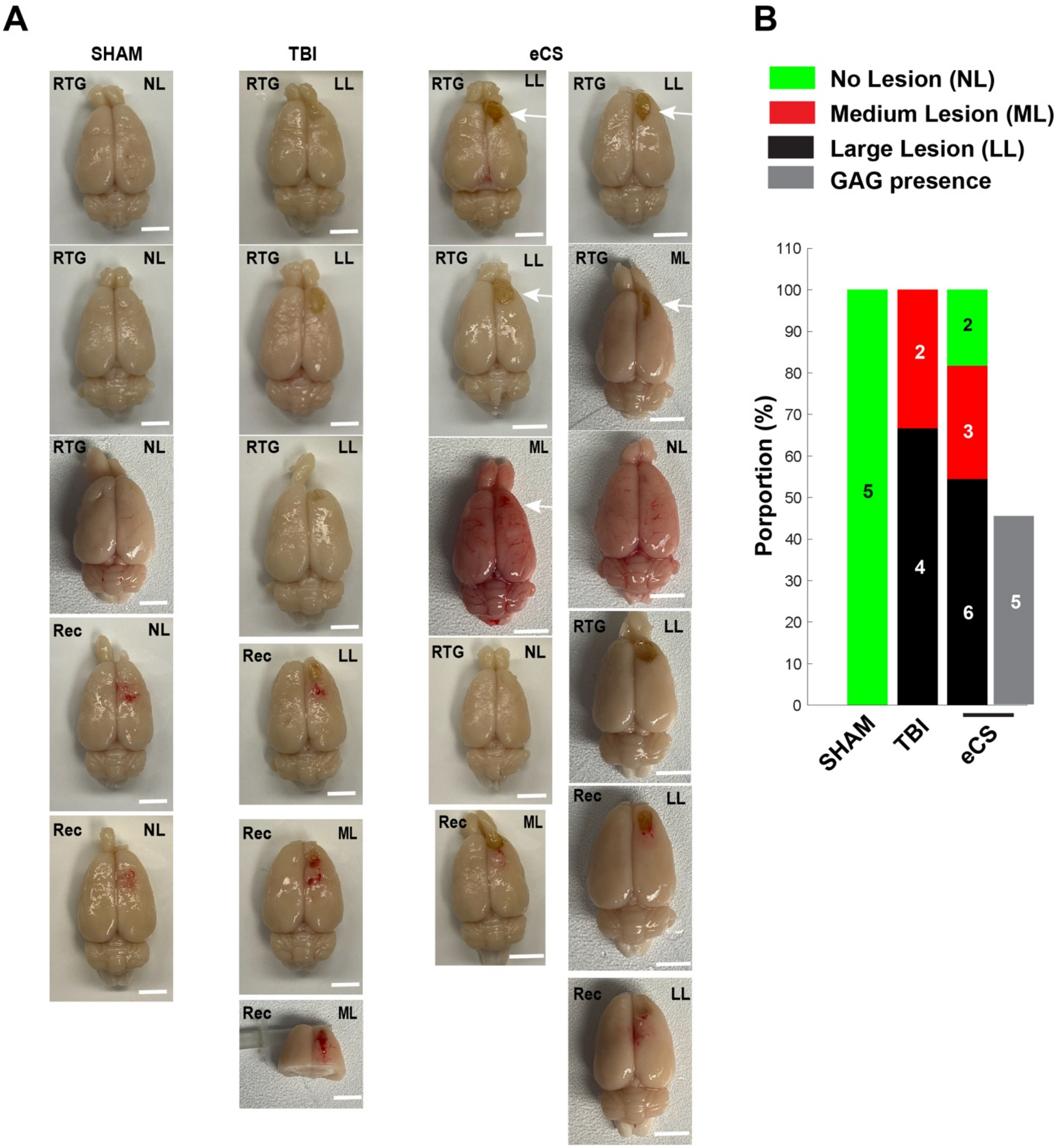
A
SHAM
TBI
eCS
RTG
Rec
NL
LL
ML
B
No Lesion (NL)
Medium Lesion (ML)
Large Lesion (LL)
GAG presence
Porportion (%)
SHAM
TBI
eCS

**Supplementary Figure 13** – Gross quantification of lesion size and eCS presence 10 weeks post-TBI

A – Top view for all rat brains used in the skilled reach task with follow up skilled reach task (SRT)-Arc or STIM-PAW-Arc assessment. White arrows indicate the observed presence of eCS-GAG implants immediately post-brain extraction. Scale bar = 2mm.

B – Cumulative distribution of lesion size, measured based on the initially delineation of the injury. In the eCS group, among the 6 Large lesion rats, 5 out of 6 showed presence of an implant at 10 weeks post-TBI (white arrow indicates eCS presence). RTG: rats were used for the reach-to-grasp assessment with immediate early gene Arc; Rec: rats were used for the 32-channel linear probe recording and paw stimulation under ketamine with immediate early gene quantification.

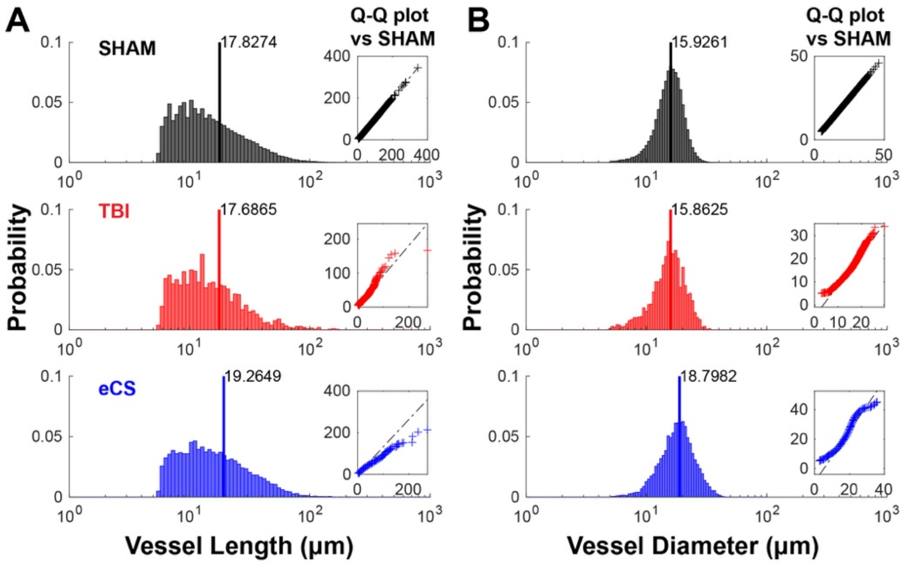


**Supplementary Figure 14** – eCS shift in blood vessel length and diameter distributions 10 weeks post-TBI in RFAa

A – Blood vessel length distribution for the SHAM (top panel), TBI (middle panel) and eCS (bottom panel) rats; data is pooled from 5 rats per groups; for each rat, the ROI volume was fixed to 500 $\mu m^3$; Inset represent the quartile-quartile plot of the panel distribution against SHAM distribution to illustrate the difference in the distribution skewness and goodness-of-fit.

B – Blood vessel diameter distribution for the SHAM (top panel), TBI (middle panel) and eCS (bottom panel) rats; data is pooled from 5 rats per groups; for each rat, the ROI volume was fixed to 500 $\mu m^3$; Inset represent the quartile-quartile plot of the panel distribution against SHAM distribution to illustrate the difference in the distribution skewness and goodness-of-fit. RFAa: anterior RFA.

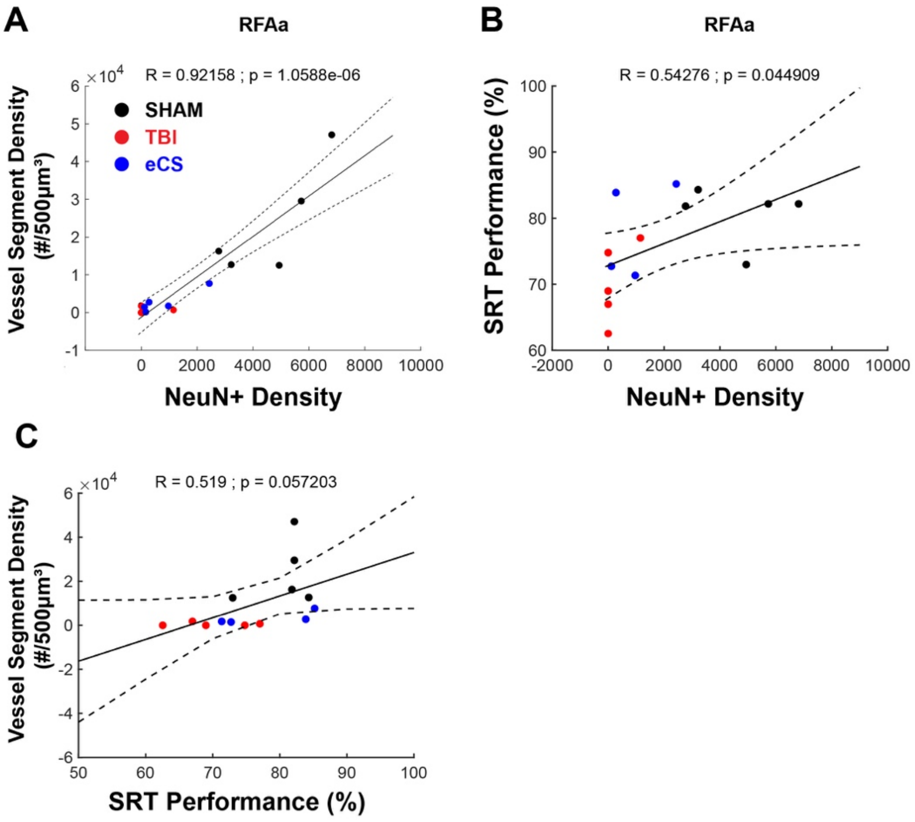


**Supplementary Figure 15**: Vascularization in RFA anterior correlates with neuronal presence and forelimb behavior performance

A – Pearson's correlation between the count of NeuN+ and blood vessel segment density within the same 500 $\mu m^3$ ROI RFAa (n = 13/15; two TBI rats showed no cellular and vasculature presence); Pearson's correlation, R = 0.92, $p < 0.001$.

B – Pearson's correlation between the count of NeuN+ in RFAa and SRT performance at the skilled reaching task (n = 13/15; two TBI rats showed no cellular presence); Pearson's correlation, R = 0.54, p = 0.045. SRT: skilled reach task.

C – Pearson's correlation between the SRT performance at the skilled reaching task and blood vessel segment density within the same 500 $\mu m^3$ ROI RFAa (n = 13/15; two TBI rats showed no vasculature presence); Pearson's correlation, R = 0.519, p = 0.057.

### Supplementary Videos

#### Supplementary Video 1: eCS implant Magnified view

https://drive.google.com/file/d/1uYce8nLLyNbqv0lGe4UOIoAOUIC1xa_C/view?usp=sharing

Volumetric imaging of a representative eCS implant at 10 weeks post-TBI. Sample was cleared using iDisco+, stained using NeuN+ (Neuronal), Reca-1 (Endothelial cells/Vasculature) and image using a Lightsheet Microscope. Magnification x4; Animation created using Imaris 6.5.

#### Supplementary Video 2: Sham prefrontal cortex over view

A – Whole frontal section, magnification x0.63: https://drive.google.com/file/d/1_BbwWFq_9_l-6DdIIZn8BcvvMuifzLKw/view?usp=sharing

B – Ipsilesional Hemisphere, Magnification x2: https://drive.google.com/file/d/1xjHH_NMeXZ2d1WCszi_nuNSFJWgPDjLV/view?usp=sharing

Volumetric imaging of a representative Sham prefrontal cortex at 10 weeks post-intervention. Sample was cleared using iDisco+, stained using NeuN+ (Neuronal), Reca-1 (Endothelial cells/Vasculature), and Arc (activity-related immediate early gene); imaged using a Lightsheet Microscope. Animation created using Imaris 6.5.

#### Supplementary Video 3: TBI prefrontal cortex over view

A – Whole frontal section, magnification x0.63: https://drive.google.com/file/d/1MSKZCGr7BBy_Y3CM45TFvSVoShcmIC9y/view?usp=sharing

B – Ipsilesional Hemisphere, Magnification x2: https://drive.google.com/file/d/16z-yqK33P7KGZqJqRa7o2-DSNGzch2HG/view?usp=sharing

Volumetric imaging of a representative TBI prefrontal cortex at 10 weeks post-TBI. Sample was cleared using iDisco+, stained using NeuN+ (Neuronal), Reca-1 (Endothelial cells/Vasculature), and Arc (activity-related immediate early gene); imaged using a Lightsheet Microscope. Magnification x0.63; Animation created using Imaris 6.5.

**Supplementary Video 4: eCS prefrontal cortex over view**

A – Whole frontal section NeuN, magnification x0.63:
https://drive.google.com/file/d/1mun49Ev0ZDWjTLZyjADUkzGPlgffh1uB/view?usp=sharing

B - Whole frontal section Reca-1, magnification x0.63:
https://drive.google.com/file/d/1Zec0BXPF14EjoSAWi9uIX4i4Eqty7vJ3/view?usp=sharing

C - Ipsilesional Hemisphere, Magnification x2:
https://drive.google.com/file/d/1l82SM38Qm1Y0VU_ydYZzSBxDZKje15dh/view?usp=sharing

Volumetric imaging of a representative eCS prefrontal cortex at 10 weeks post-TBI/eCS implant. Sample was cleared using iDisco+, stained using NeuN+ (A - Neuronal) and Reca-1 (B - Endothelial cells/Vasculature); imaged using a Lightsheet Microscope. Animation created using Imaris 6.5.
